# Lipid accumulation promotes scission of caveolae

**DOI:** 10.1101/2020.01.22.915173

**Authors:** Madlen Hubert, Elin Larsson, Naga Venkata Gayathri Vegesna, Maria Ahnlund, Annika I. Johansson, Lindon W. K. Moodie, Richard Lundmark

**Affiliations:** Department of Integrative Medical Biology, Umeå University, SE-901 87 Umeå, Sweden; Swedish Metabolomics Centre, Department of Forest Genetics and Plant Physiology, Swedish University of Agricultural Sciences, SE-901 83 Umeå, Sweden; Swedish Metabolomics Centre, Department of Molecular Biology, Umeå University, SE-901 83 Umeå, Sweden; Department of Chemistry, Umeå University, SE-901 87 Umeå, Sweden

**Keywords:** caveolae, EHD2, cell surface stability, scission, fusogenic liposomes, glycosphingolipids, cholesterol, membrane lipid composition, single particle tracking, correlative light electron microscopy

## Abstract

Caveolae, bulb-shaped invaginations of the plasma membrane (PM), show distinct behaviors of scission and fusion at the cell surface. Although it is known that caveolae are enriched in cholesterol and sphingolipids, exactly how lipid composition influences caveolae surface stability has not yet been elucidated. Accordingly, we inserted specific lipids into the PM of cells via membrane fusion and studied acute effects on caveolae dynamics. We demonstrate that cholesterol and glycosphingolipids specifically accumulate in caveolae, which decreases their neck diameter and drives their scission from the cell surface. The lipid-induced scission was counteracted by the ATPase EHD2. We propose that lipid accumulation in caveolae generates an intrinsically unstable domain prone to scission if not balanced by the restraining force of EHD2 at the neck. Our work advances the understanding of how lipids contribute to caveolae dynamics, providing a mechanistic link between caveolae and their ability to sense the PM lipid composition.

**SUMMARY:** Caveolae serve as mechanoprotectors and membrane buffers but their specific role in sensing plasma membrane lipid composition remains unclear. Hubert et al. show that cholesterol and glycosphingolipids accumulate in caveolae and drive subsequent scission from the cell surface. These results provide new insight into how lipids contribute to budding and scission of membrane domains in cells.

## INTRODUCTION

Caveolae are bulb-shaped invaginations of the plasma membrane (PM), enriched in cholesterol (Chol), sphingolipids and the integral membrane protein caveolin1 (Cav1) (Parton & del Pozo, 2013). Caveolae are present in most cell types, with a particularly high density in endothelial cells, adipocytes and smooth muscle cells. Absence or malfunction of caveolae is associated with a number of conditions such as lipodystrophy, muscular dystrophy and cardiovascular diseases (Cohen et al., 2004; Pilch & Liu, 2011). Whilst the mechanism of how caveolae dysregulation drives the phenotype of disease is not well understood, they have been proposed to serve as signaling platforms, endocytic carriers, and PM reservoirs involved in mechanoprotective processes or lipid buffering (Parton & del Pozo, 2013; Sinha et al., 2011). In adipocytes, which are key lipid homeostasis regulators, caveolae are estimated to account for more than 50% of the surface area (Thorn et al., 2003). The clinical manifestation of caveolae loss in both patients (Cao et al., 2008; Hayashi et al., 2009; Kim et al., 2008) and mouse models (Liu et al., 2008; Razani et al., 2002) reveals severe malfunction of adipocytes, in addition to other cell types involved in lipid turnover and storage.

Biogenesis of caveolae is tightly coupled to the PM lipid composition and is thought to be driven by Chol-sensitive oligomerization of Cav1 and subsequent association with the cavin coat proteins (Fig. S1A) (Parton & del Pozo, 2013). Cav1 is embedded into the lipid bilayer via its intramembrane and scaffolding domains, which interact with Chol (Parton & del Pozo, 2013). Chol depletion from the PM causes caveolae disassembly, leading to the disassociation of cavins from Cav1, which then disseminates throughout the PM (Morén et al., 2012; Rothberg et al., 1992). Relative to the bulk PM, phosphatidylserine, glycosphingolipid (GSL) and sphingomyelin (SM) has also been proposed to be enriched in caveolae (Hirama et al., 2017; Örtegren et al., 2004; Singh et al., 2010). However, it has not been determined whether specific lipids are enriched in caveolae in living cells and, furthermore, how exactly they influence biogenesis, *i.e.*, through a general physical effect on the membrane bilayer or via direct interactions with the caveolae coat proteins. Additionally, it is not known whether these lipids can diffuse freely in and out of the caveolae bulb or if they are sequestered by interactions with the caveolae coat and membrane curvature restraints.

Whilst caveolae are typically associated with the PM as bulb-shaped invaginations, they also exhibit dynamic behaviour including flattening (Nassoy & Lamaze, 2012), short-range cycles of fission and fusion with the PM, and endocytosis (Boucrot et al., 2011; Morén et al., 2012; Pelkmans & Zerial, 2005). Caveolae are stabilized at the cell surface by the ATPase Eps-15 homology domain-containing protein 2 (EHD2), which oligomerizes around the neck of caveolae, restraining their scission (Fig. S1A) (Morén et al., 2012; Stoeber et al., 2012). EHD2 extensively colocalizes with Cav1 and most of the membrane-associated EHD2 is found in caveolae (Morén et al., 2012). Although not considered part of the caveolae coat, EHD2 appears to be a critical component for maintaining caveolae integrity in terms of surface attachment. Despite that all the above-mentioned proposed functions of caveolae would heavily depend on whether they are surface associated or released, it is not clear how the balance between these states is controlled physiologically. Furthermore, the biological function of their atypical dynamics remains elusive.

Lipids are also thought to influence caveolae dynamics, *e.g.*, addition of bovine serum albumin-complexed lactosyl ceramide (LacCer) and elevated levels of Chol have been proposed to reduce the number of surface-connected caveolae and increase their mobility (Le Lay et al., 2006; Sharma et al., 2004). However, despite their essential structural role in caveolae, little is known about how they influence caveolae biogenesis and dynamics. This knowledge gap can be partly attributed to limitations associated with the current methods employed to study such phenomena. Drugs such as statins, that inhibit Chol synthesis, require multi-day treatments, and, in addition to altering transcriptional regulation, may also elicit major secondary effects (Crescencio et al., 2009). This results in downregulated expression of Cav1, making it difficult to decipher between the effects of Chol levels on caveolae biogenesis and caveolae dynamics. In addition, drugs such as myriocin, which inhibit sphingosine synthesis, will also affect the levels of all sphingolipid species, thus hampering direct conclusions. BSA-coupled Bodipy-LacCer has been used as a fluorescent marker of endocytosis (Puri et al., 2001; Sharma et al., 2004; Singh et al., 2006; Singh et al., 2003), but this procedure involved PM loading at 4°C followed by a temperature shift to 37°C, known to heavily influence membrane fluidity and exacerbate endocytosis (Kleusch et al., 2012). While previous work indicates that lipids might influence both caveolae numbers and their dynamics, they have been unable to address whether caveolae dynamics respond directly to alterations in PM lipid composition and if proposed effects are dependent on concentration or species of different lipids present. In general, our understanding of the levels of quantitative changes in PM lipid composition that can be sensed and controlled is relatively sparse, not to mention the alteration in lipid composition required to influence caveolae. It is also not known if lipids could affect caveolae dynamics by changing the composition of the caveolae bulb or the surrounding membrane and whether the proposed effects on caveolae mobility is caused by direct effects on caveolae scission from the cell surface.

To address this, we aimed to rapidly and selectively manipulate cellular membrane lipid composition in a system where both the lipids and caveolae could be tracked. Here, we have applied fusogenic liposomes that allowed us to directly insert specific unlabeled or fluorescently-labeled lipids into the PM of living cells and study their effect on caveolae dynamics. Our data shows that a relatively small increase in glycosphingolipids and Chol results in their accumulation in caveolae, reducing the caveolae neck diameter, and driving caveolae scission from the PM. EHD2 was identified to counterbalance the stability of caveolae in response to lipid composition and in accordance with a recent study (Matthäus et al., 2019; Morén et al., 2019), we describe a key regulatory role of EHD2 in lipid homeostasis.

## RESULTS

### Lipids rapidly insert into the PM of living cells via liposome-mediated membrane fusion

As a tool to study the effects of an altered lipid composition on caveolae dynamics, we employed fusogenic liposomes. This enabled us to rapidly insert lipids into the PM of HeLa cells via membrane fusion (Fig. 1A). To assess the effect of lipids known to be enriched in caveolae, different Bodipy-labeled analogues of sphingolipids (Cer, SM C_5_ and SM C_12_), GSLs [ganglioside GM1 and lactosyl ceramide (LacCer)], Chol and phosphatidyl ethanolamine (PE) (Fig. S1B) were incorporated into liposomes [DOPE/Dotap/Bodipy-tagged lipid (47.5/47.5/5)] (Csiszar et al., 2010). Liposomes had diameters between 160-300 nm (Fig. S1C) and an average fluorescence per liposome of 600 a.u (Fig. S1D). The Bodipy fluorophore allowed us to track and quantify the lipid incorporation in the PM and study colocalization with caveolae components. To ensure the observed effects are not significantly influenced by the fluorophore motif, the results were verified with unlabeled lipids. Fusion of the liposomes with the PM of HeLa cells occurred immediately upon contact and the lipids were rapidly distributed throughout the basal membrane, as observed using live-cell TIRF microscopy (Figs. 1B and S1E, exemplified by LacCer). The total fluorescence attributed to the Bodipy motif increased uniformly in various regions of interest (ROIs) (Fig. 1B, bar plot).

**Fig. 1.**
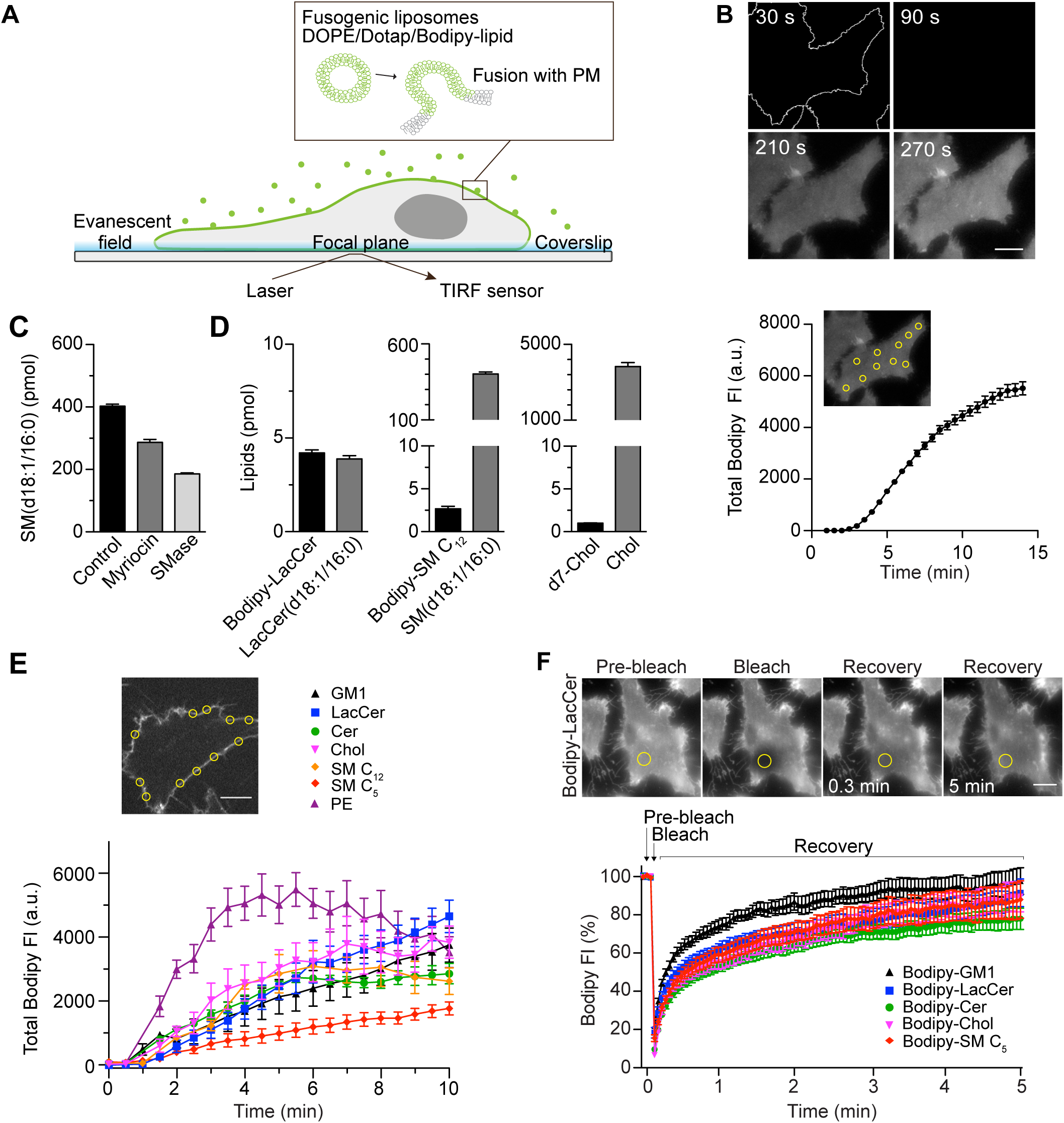
Rapid insertion of Bodipy-labeled lipids into the PM of living cells using fusogenic liposomes. **(A)** Fusogenic liposomes are used to insert Bodipy-labeled lipids into the PM. Their rapid distribution is followed in real time using TIRFM. **(B)** Image sequence of Bodipy-LacCer distribution throughout basal membrane of HeLa cells. Total Bodipy fluorescence intensity (FI) was measured within ROIs (yellow insert) using Zeiss Zen interface. *n* = 10, three independent experiments, mean ± SEM. **(C)** Quantification of endogenous SM(d18:1/16:0) using LC-ESI-MS/MS in untreated control cells or cells treated with SMase or myriocin for 2 h or 24 h, respectively. Data are shown as mean + SD. **(D)** Quantification of Bodipy- or d7-labeled lipids (black bars) and endogenous lipids (grey bars) in cells following incubation of cells with fusogenic liposomes. Analysis was performed using mass spectrometry. Data are shown as mean + SD. **(E)** Incorporation rate of Bodipy-lipids into PM of live cells. HeLa cells were treated with fusogenic liposomes (final total lipid concentration 7 nmol/mL). Total Bodipy fluorescence intensity (FI) was measured within circular ROIs (see insert) in a confocal section using spinning disk microscopy. Ten ROIs were analyzed using the Zeiss Zen system software. *n* ≥ 2, two independent experiments, mean ± SEM. Scale bars, 10 μm. **(F)** TIRF FRAP of Bodipy-lipids after incorporation into PM of HeLa cells. A circular ROI was photobleached and recovery of Bodipy FI was monitored over 5 min. Bodipy FI was normalized to background and reference. *n* ≥ 10, mean ± SEM.

To determine the amounts of lipids that were incorporated in the membrane through liposome fusion, we used quantitative mass spectrometry on whole cells as 90% of these lipids are located in the PM (Lorizate et al., 2013). The method was verified by altering the lipid composition using myriocin (24h treatment) or sphingomyelinase (SMase, 2h treatment), which are known to lower the levels of sphingomyelin (Gulshan et al., 2013). Analysis showed that these treatments drastically decreased SM(d18:1/16:0) levels, the major endogenous species of SM (Fig. 1C). Next, we incubated cells with fusogenic liposomes containing Bodipy-labeled LacCer or SM C_12_, and analyzed the lipid composition by LC-ESI-MS/MS. The detected levels of endogenous LacCer(d18:1/16:0) and SM(d18:1/16:0) in untreated control samples were in agreement with previously reported levels in HeLa cells (Kjellberg et al., 2014). In comparison, in samples treated with fusogenic liposomes, the incorporated Bodipy-lipids could be detected (Figs. S1F-G). The incorporated levels of Bodipy-LacCer and Bodipy-SM C_12_ per 400 000 cells were measured to be 4.2 pmol and 2.7 pmol, respectively, (*i.e.*, 6.3 × 10^6^ and 4.0 × 10^6^ lipids/cell) (Fig. 1D). To assess the incorporation efficiency of Chol, deuterium labeled Chol, d7-Chol, was included into fusogenic liposomes. GC-MS/MS analysis revealed that d7-Chol was incorporated to similar levels as Bodipy-labeled LacCer and SM C_12_. Given that the PM of these HeLa cells harbor around 7 × 10^9^ lipids/cell (see Methods section for details), the levels of Bodipy- and d7-labeled lipids detected by mass spectrometry led to a 0.02-0.09% increase in specific labeled lipids and a 0.4-1.6% of total lipids.

To determine the rate of incorporation of the different Bodipy-labeled lipid species, we used spinning disk microscopy in a central confocal plane of the cell (Fig. 1E). Quantitative analysis of lipid incorporation into the PM over time revealed similar levels for most lipids ranging from 1900 to 4800 arbitrary units at 10 min (Fig. 1E). The variation in incorporation rates between the different lipids may be due to marginal differences in the fusogenicity of the liposomes or differences in the PM-turnover of each particular lipid. To monitor the lateral diffusion of the Bodipy-lipids within the PM, cells were incubated with fusogenic liposomes for 10 min, and the recovery of Bodipy fluorescence in a defined region of interest (ROI) after bleaching was monitored. Lateral diffusion within the PM was similar between the different Bodipy-lipids employed (Fig. 1F).

To conclude, the use of fusogenic liposomes enabled rapid incorporation of approximately 4 × 10^6^ specific lipids into the PM per living cell over 10 minutes. Based on our tracking data, we estimated that each cell contains around 300 caveolae, comprising around 0.1% of the surface area. Caveolae are approximately 60 nm in diameter, and each lipid occupies 0.62 Å. Therefore *ca.* 10 × 10^6^ lipids are contained within the caveolae, of which 50% is Chol. The amount of specific incorporated lipids in our system is therefore about half of the total amount of lipids contained within caveolae. The immediate addition of extra lipids to the PM did not result in a detectable effect on the cell volume (Fig. S1H).

### GSLs and Chol decrease the surface stability of caveolae

We next aimed to elucidate if lipids are involved in controlling the balance between stable and dynamic caveolae at the PM and if effects could be attributed to individual lipid species. To visualize caveolae, we generated a stable mammalian Flp-In T-Rex HeLa cell line expressing Cav1-mCherry, hereafter named Cav1-mCh HeLa cells. Protein expression was induced by doxycycline (Dox) to achieve expression of Cav1-mCherry at similar level as endogenous Cav1 (Fig. S1I). Using TIRF microscopy and single-particle tracking, we determined the time each Cav1-mCh positive punctuate structure spent at the PM (track duration) and the speed of an object (track mean speed) in, or close to, the PM (see Method section for detailed tracking parameters). Given the previously reported surface dynamics of caveolae (Boucrot et al., 2011; Mohan et al., 2015; Pelkmans & Zerial, 2005), we postulated that stable caveolae will have a long duration and low speed, limited by their lateral diffusion in the PM (Fig. 2A, “Stable”). Caveolae that scission off (“Scissioned intermediate”) or re-fuse (“Fused intermediate”) with the PM during the recording period will have an intermediate duration and moderate speed, whereas caveolae that undergo rounds of scission and fusion (“Surface adjacent”), but remain close to the surface will have a high speed and short duration as they are not stably fused with the PM and short duration (Fig. 2A). We did indeed observe a clear correlation between the track duration and track mean speed where, in general, short tracks exhibited higher speeds, whereas long tracks displayed lower speeds (Figs. 2B and S2A). Although the numbers of caveolae in each cell were similar at the beginning and end of the recording, we found that the number of tracks far exceeded the caveolae numbers (Fig. S2C). This was expected as surface adjacent caveolae would give rise to several tracks. However, a drop in the fluorescent signal just below the set threshold value, would also contribute to a divided track resulting in an overestimation of short tracks versus long tracks. Therefore, we did not consider the average durations as absolute, but rather used them to compare between experimental runs with differing conditions. To verify that the tracking was sensitive to differences in caveolae dynamics, we depleted cells of EHD2, which has been shown to stabilize caveolae to the cell surface (Morén et al., 2012; Stoeber et al., 2012). Particle tracking analysis showed that the pool of surface adjacent caveolae increased, while the pool of stable caveolae decreased (Figs. 2B’ and S2A’). When the average track duration was considered, this translated into a 0.65 fold change compared to control cells (Fig. 3E), proving that the particle tracking indeed was sensitive enough to register caveolae dynamic changes.

**Fig. 2.**
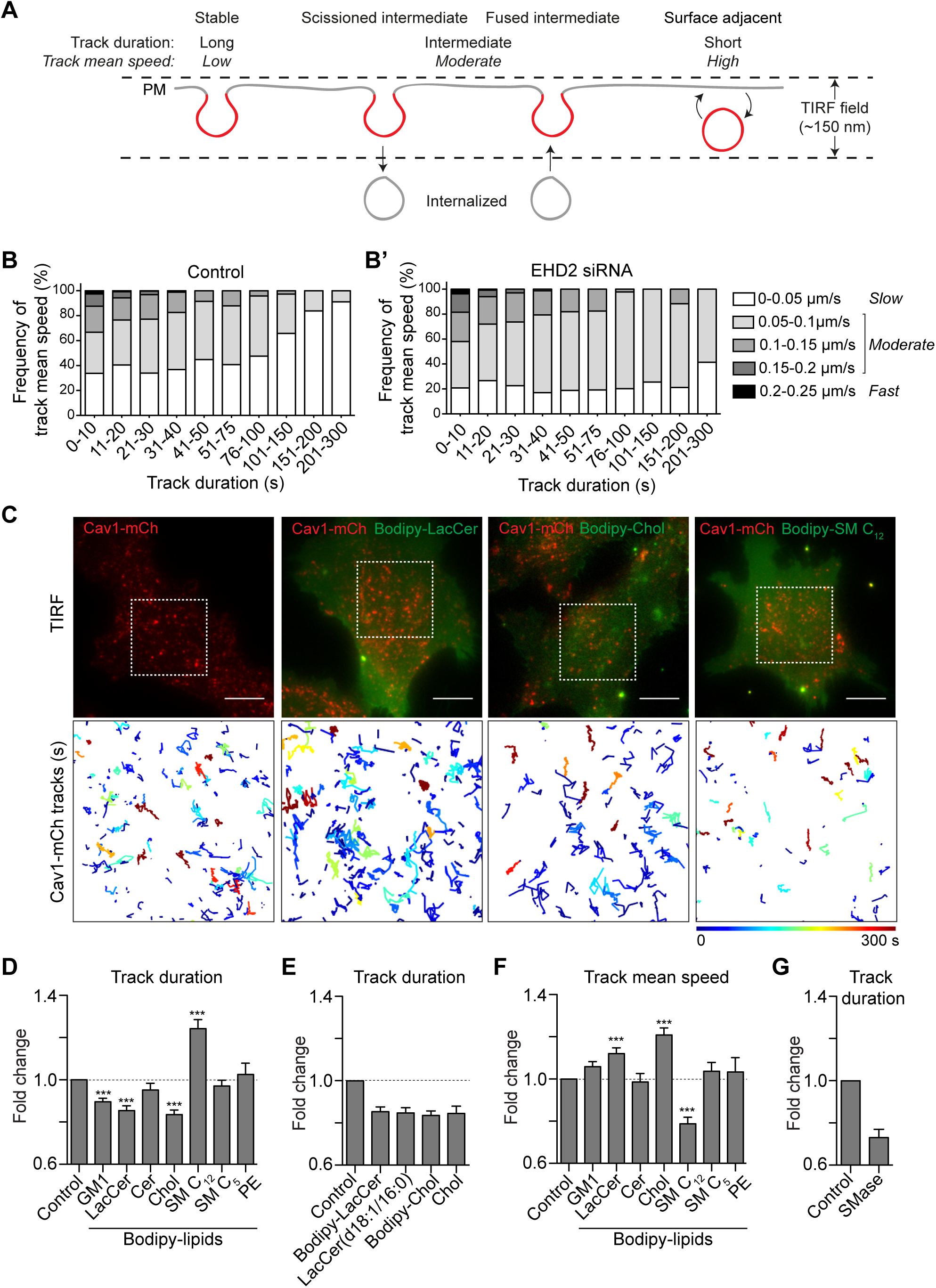
GSLs and Chol decrease the surface stability of caveolae. **(A)** Scheme showing different dynamic behaviors of caveolae. **(B, B’)** Distribution of track mean speed amongst subpopulations of track duration of Cav1-mCh structures **(B)** and after EHD2 depletion **(B’)**. Five datasets for each condition were analyzed from TIRF live cell movies. **(C)** Representative images from TIRF movies of Cav1-mCh HeLa cells and after 15 min incubation with liposomes containing Bodipy-lipids. Color-coded trajectories illustrate time that structures can be tracked at PM over 5 min (dotted square). Scale bars, 10 μm. See Video 1-4. **(D-E)** Quantification of track duration of Cav1-mCh structures from TIRF movies after incubation with liposomes containing labeled **(D)** or unlabeled lipids **(E)**. Fold changes are relative to control (Cav1-mCh). (D) *n* ≥ 8, at least two independent experiments; (E) *n* ≥ 8, two independent experiments, ***, *P* ≤ 0.001 vs. control. **(F)** Quantification of track mean speed of Cav1-mCh structures from TIRF movies (same cells as in D). **(G)** Quantification of track duration of Cav1-mCh structures from TIRF movies following incubation with SMase for 2 h. Fold changes are relative to control (Cav1-mCh). *n* ≥ 5. All analyses were performed using Imaris software and data are shown as mean + SEM.

**Fig. 3.**
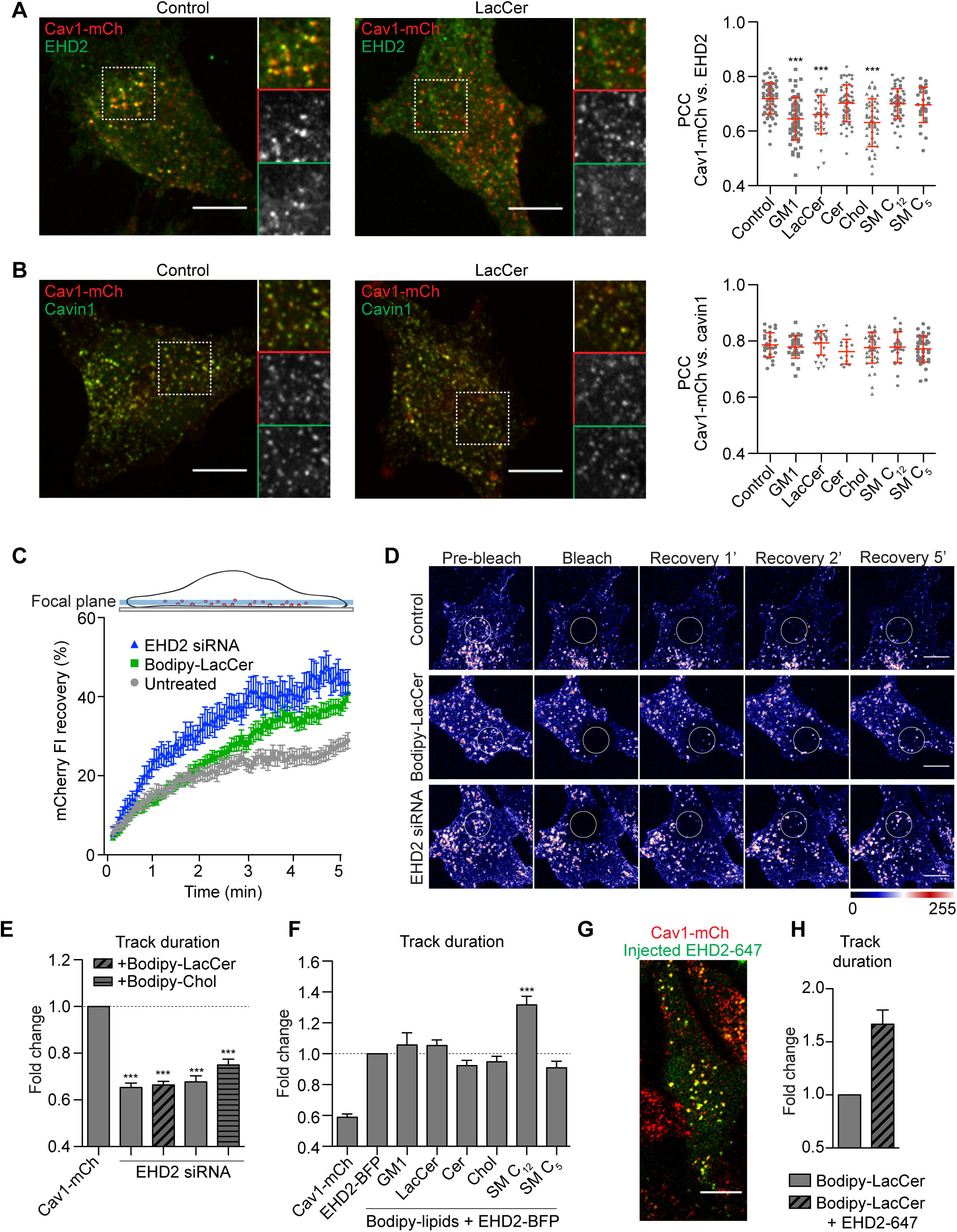
Chol and GSLs induce surface release of caveolae via an EHD2-dependent mechanism. **(A)** Representative images of maximum projected confocal z-stacks of Cav1-mCh HeLa cells. Untreated cells or cells treated with LacCer-Bodipy liposomes for 1 h, fixed and immunostained for endogenous EHD2. High-magnification images (dotted square) show localization of EHD2 to Cav1-mCh (see scatterplot for quantification). *n* ≥ 60, two independent experiments, mean ± SEM. ***, *P* ≤ 0.001 vs. control. **(B)** Experimental protocols analogous to (A), with exception of endogenous cavin1 immunostaining. *n* ≥ 60, mean ± SEM. **(C)** Confocal FRAP of Cav1-mCh HeLa cells treated with either EHD2 siRNA or Bodipy-LacCer liposomes. A ROI was photobleached and recovery of mCherry FI monitored over 5 min. mCherry FI was normalized to background and reference. *n* ≥ 10, mean ± SEM. **(D)** Representative time-lapse series showing control Cav1-mCh HeLa cells and cells treated with either EHD2 siRNA or Bodipy-LacCer liposomes. The photobleached area is outlined with white circles. mCherry FI is intensity coded using LUT. **(E)** Effects of lipids on track duration of Cav1-mCh structures were analyzed following siRNA-mediated depletion of EHD2. *n* ≥ 8, two independent experiments, mean + SEM. **(F)** Quantification of track duration of Cav1-mCh HeLa cells transiently expressing EHD2-BFP with or without incubation with liposomes. Changes in track duration are relative to control (indicated by dotted line). *n* ≥ 8, two independent experiments, mean + SEM. ***, *P* ≤ 0.001 vs. control cells. **(G)** Representative live cell confocal image of EHD2-647 microinjected into Cav1-mCh HeLa cells. (**H**) Quantification of track duration of Cav1-mCh cells treated with Bodipy-LacCer and following microinjection of EHD2-647. *n* = 8, mean + SEM. All scale bars, 10 μm.

Next, we screened the influence of different lipid species on caveolae mobility at the PM using tracking. To do this, fusogenic liposomes loaded with relevant lipids were added to Cav1-mCh HeLa cells and TIRF movies were recorded immediately (Fig. 2C). PE was used in control liposomes as it is abundant in the PM. Following incorporation of PE, caveolae dynamics remained unchanged, showing that the fusion of liposomes did not obstruct caveolae dynamics (Figs. 2D and S2B). In comparison to controls, Bodipy-labeled GSLs (GM1 and LacCer) and Chol significantly reduced the lifetime of caveolae at the cell surface as indicated by decreased track duration measurements (Figs. 2C-D and S2B, Video 1-3). Besides enhanced mobility, caveolae also showed an increase in mean speed (Fig. 2E). For example, the treatment with LacCer gave a ratio of surface adjacent caveolae versus stable that was comparable to the EHD2 depletion (Figs. 2B’, S2A’ and S2F-G). A direct comparison between LacCer and Cer revealed that Cer did not enhance caveolae dynamics in a similar fashion (Figs. 2D and S2B). No difference in the number of caveolae present in the PM was observed before and after the addition of liposomes (Fig. S2D). To verify that the effect was not due to the the Bodipy label, we treated cells with liposomes containing either unlabeled LacCer [(LacCer(d18:1/16:0)] or unlabeled Chol and quantified the track duration. This showed that the unlabeled lipids had the same effect on the track duration as the corresponding Bodipy-labeled analogues (Fig. 2F).

When cells were treated with Bodipy-SM C_12_, most of the caveolae were stable at the PM (Fig. 2C-D). This was characterized by a dramatic increase in track duration, and a reduction of the track mean speed (Figs. 2D and 2E). To further investigate the role of SM, we analyzed caveolae duration following SMase treatment, and found that this resulted in a decreased track duration, in agreement with a surface-stabilizing role for this lipid (Fig. 2G).

### Chol and GSLs induce surface release of caveolae via an EHD2-dependent mechanism

EHD2 normally localizes with the majority of surface associated caveolae (Morén et al., 2012). We aimed to address if the increased caveolae dynamics induced by either Chol or GSLs were due to their PM release, as characterized by loss of the stabilizing protein EHD2. Therefore, we treated Cav1-mCh HeLa cells with the different fusogenic liposomes and visualized endogenous EHD2 using indirect immunofluorescent labeling (Fig. 3A). These experiments revealed that incorporation of GM1, LacCer or Chol into the PM led to a significantly lower amount of EHD2 localized to Cav1 (Fig. 3A, scatter plot). This data suggests that the caveolae release induced by increased PM levels of LacCer and Chol is due to a loss of EHD2-mediated stabilization. Conversely, Cer and SM C_12_, as well as its short chain analogue SM C_5_, did not appear to have any significant effect on the association of EHD2 with Cav1 (Fig. 3A, scatter plot). Further experiments showed that, following lipid treatment, the majority of caveolae remained associated with cavin1, revealing that no disruption of the caveolae coat, and subsequent release of cavin1, occurred (Fig. 3B).

Since increased scission of caveolae from the cell surface results in more mobile intracellular caveolar vesicles (Stoeber et al., 2012), we performed fluorescence recovery after photobleaching (FRAP) experiments to investigate the recovery rate of caveolae. Addition of LacCer resulted in more mobile caveolae inside the cells in comparison with control cells (Figs. 3C-D). The recovery rate after LacCer addition was similar to the rate in EHD2-depleted cells (Figs. 3C-D). This further supported the hypothesis that lipid incorporation drives caveolae scission (Figs. 3C-D). When LacCer or Chol was added to EHD2-depleted cells we found that the caveolae track duration was not further reduced in comparison with EHD2-depleted cells (Figs. 3E and S3A-B). These results supported our hypothesis that the lipid-induced effect on caveolae dynamics was due to the loss of surface stability mediated by EHD2, and implied that EHD2 controls the stability of caveolae in response to lipid composition. To test whether increased levels of EHD2 could restore caveolae stability after lipid treatment, we transiently expressed BFP-tagged EHD2 in Cav1-mCh HeLa cells. Analysis of TIRF live cell movies showed that EHD2-BFP positive caveolae were highly stable compared to controls (Fig. 3F). In the presence of EHD2-BFP, the destabilizing effect seen for GM1, LacCer and Chol was abolished, as demonstrated by negligible changes in track duration compared to control conditions (Fig. 3F). This suggested that excess levels of EHD2 were capable of restricting the effect of excess Chol and GSLs. In addition, tracking of caveolae in cells stably expressing Cav1-mCh and EHD2-BFP showed that most of the caveolae were positive for EHD-BFP (96%) and that this population was more stable than the population lacking EHD2-BFP (4%) (Figs. S3C-E). Surprisingly, SM C_12_ had an additive effect and led to predominantly stable caveolae (Fig. 3F), implying that the increased cell surface stability of caveolae may be due to changes in membrane fluidity rather than EHD2 recruitment.

To test if the suppression of the lipid effect by increased levels of EHD2 relied on multiple rounds of assembly and disassembly of EHD2 at caveolae, we overexpressed a BFP-tagged ATP-cycle mutant, EHD2-I157Q. The increased ATP hydrolysis rate of this mutant leads to stable association of EHD2-I157Q to caveolae and a slower exchange rate (Fig. S3F) (Daumke et al., 2007; Hoernke et al., 2017; Stoeber et al., 2012). We observed that, when co-expressed in Cav1-mCh cells, both EHD2-I157Q and EHD2 stabilized caveolae at the PM to similar extents, independent of treatment with either LacCer or Chol (Fig. S3G). This verified that stable assembly, but not disassembly of EHD2, is necessary to stabilize caveolae.

To clarify whether, in order to have a stabilizing role, EHD2 had to be caveolae-associated prior to lipid addition, fluorescently labeled, purified EHD2 (EHD2-647) was microinjected into Cav1-mCh HeLa cells (Fig. 3G). Within 20 min, EHD2-647 colocalized with Cav1, confirming that the microinjected protein was indeed recruited to caveolae (Figs. S3H-I). Next, we tested if an acute injection of EHD2-647 could rescue the effect on caveolae dynamics caused by LacCer. Strikingly, we found that exogenously added EHD2 stabilized the caveolae to the same extent as the overexpressed EHD2, demonstrating that increased levels of EHD2 can acutely reverse the increased mobility of caveolae induced by lipids (Fig. 3H).

### LacCer and Chol accumulate in caveolae and Chol is sequestered within these domains

As GSLs and Chol increased the surface release of caveolae, we aimed to determine whether there was a differential accumulation of lipids within caveolae at the PM. We treated Cav1-mCh HeLa cells with fusogenic liposomes and followed the distribution of Bodipy-labeled LacCer or Chol using live-cell TIRF microscopy. After 15 min, both lipids were found to colocalize with Cav1-mCh positive structures (Figs. 4A and S4A, Video 5-6). Data analysis was hindered by high caveolae mobility following lipid addition, and the extent of colocalization could not be quantified. To circumvent this, we overexpressed EHD2-BFP to stabilize caveolae at the PM. Interestingly, nearly 80% of caveolae positive for EHD2 were also positive for LacCer or Chol (Figs. 4B-B’ and S4B, Video 7-8). In comparison, Cer, which had no effect on caveolae dynamics, did not localize to caveolae, even in the presence of EHD2-BFP (Figs. S4C-D).

**Fig. 4.**
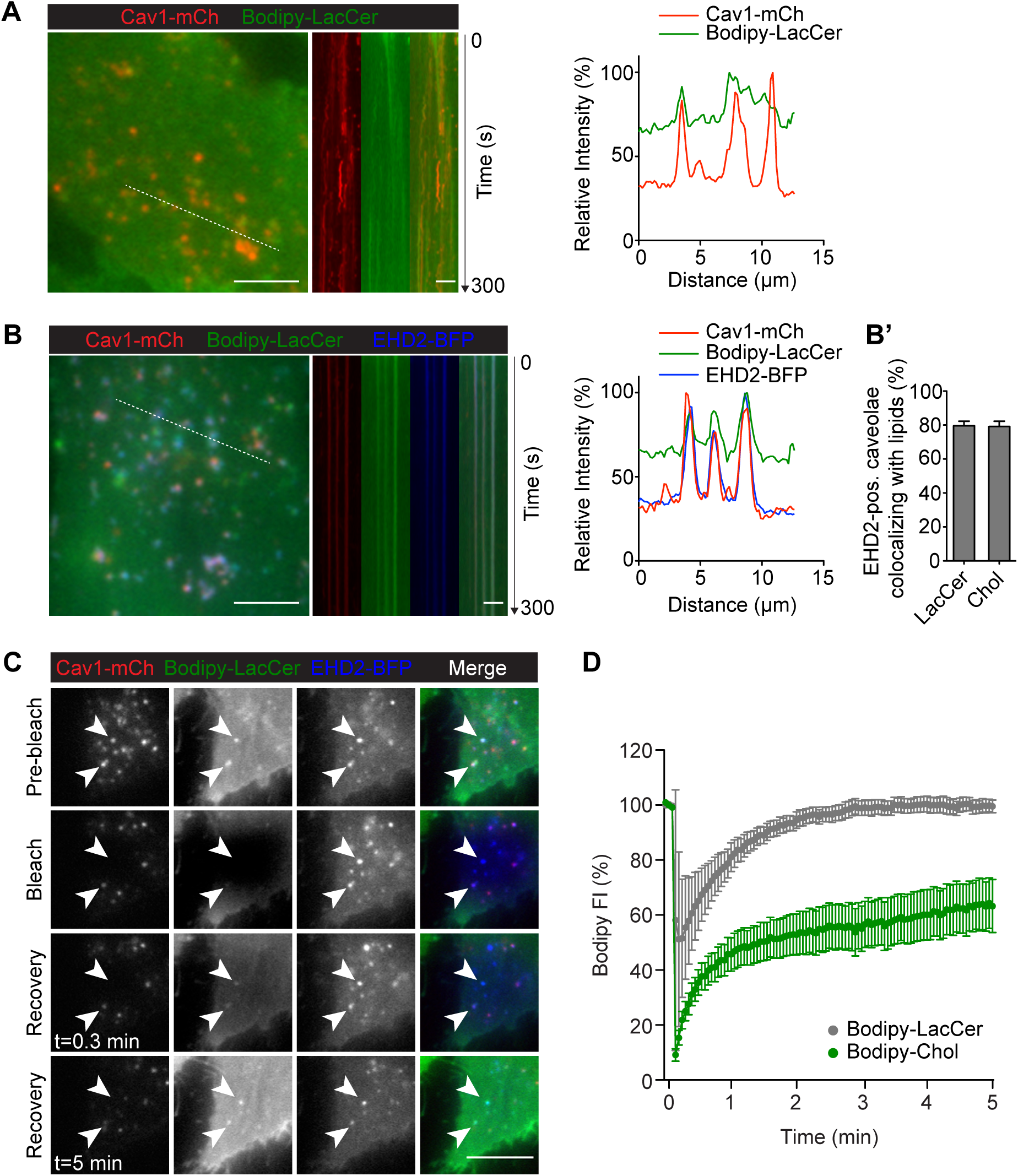
LacCer and Chol accumulate in caveolae and Chol is sequestered within these domains. **(A, B)** Cav1-mCh HeLa cells **(A)** and Cav1-mCh HeLa cells transiently expressing EHD2-BFP **(B)** were incubated with Bodipy-LacCer liposomes. White lines indicate location of kymograph and the corresponding intensity profiles illustrate localization of Bodipy-LacCer to Cav1-mCh either alone or in presence of EHD2-BFP. Intensity profiles are relative to maximum values for each sample. Scale bars, 10 μm; kymograph scale bars, 5 μm. See Video S3 and S4. **(B’)** Quantification of EHD2-positive caveolae colocalizing with lipids. *n* ≥ 8, at least two independent experiments, mean + SEM. **(C)** Cav1-mCh HeLa cells transiently expressing EHD2-BFP were incubated with Bodipy-lipids for 10 min. Following photobleaching, recovery of Bodipy signal within caveolae was monitored over time. White arrows highlight surface connected caveolae with accumulated Bodipy-LacCer. Scale bar, 5 μm. **(D)** Recovery curves of Bodipy intensities within bleached membrane ROI. Bodipy FI was normalized to background and reference. *n* ≥ 10, mean ± SEM.

To investigate the exchange of lipids between the stable caveolae and the surrounding PM, we performed FRAP experiments. The Bodipy-LacCer signal reappeared rapidly at precisely the bleached spot positive for Cav1-mCh, with a close to quantitative fluorescence recovery (Figs. 4C-D). This indicated that the lipid diffused freely throughout the PM and, following photobleaching, re-accumulated quickly within caveolae. In comparison, Bodipy-Chol recovered much slower with 60% of the initial signal being restored after 5 min acquisition time (Figs. 4D and S4E). This showed that there is a large immobile pool of Chol in caveolae that is sequestered from the rest of the PM. Our data suggests that both LacCer and Chol are highly enriched in caveolae and, while the lateral diffusion of LacCer in and out of caveolae is high, Chol is restrained to this invagination.

### Chol accumulation reduces the caveolae diameter in 3T3-L1 adipocytes

To elucidate whether lipid accumulation affected the overall morphology of surface connected caveolae in Cav1-mCh HeLa cells, we analyzed their ultrastructure in cells overexpressing EHD2. Since the number of caveolae in the PM of these cells is relatively low, we used correlative light electron microscopy (CLEM) to specifically identify fluorescently tagged caveolae by combining light microscopy with the higher resolution images of transmission electron microscopy (TEM). Fluorescence light microscopy images of Cav1-mCh HeLa cells were superimposed with correlative electron micrographs to find the closest match of fluorescence signal to structure using the nuclear stain as a guide (Figs. 5A and S5A). The surface connected caveolae in cells treated with LacCer and Chol, displayed a similar flask-shaped morphology as seen in control cells, verifying that lipid addition did not majorly distort caveolae morphology. To be able to quantitatively assess differences in morphology, we differentiated 3T3-L1 cells to adipocytes, which results in upregulation of Cav1 and EHD2 (Fig. S5B) (Morén et al., 2019), and formation of a large number of caveolae (Thorn et al., 2003) that could be clearly distinguished from clathrin-coated pits (Fig. S5C). These cells also provided a more physiologically relevant system since adipocytes are the main source of cholesterol storage and efflux (Krause & Hartman, 1984). Quantification of lipid incorporation verified that fusogenic liposomes could be used to insert specific lipids into the PM of these cells (Fig. S5D). Using TEM, we analyzed the dimensions of caveolae before and after Chol addition (Figs. 5B-E). We found that the neck diameter of surface associated caveolae were significantly decreased and more homogeneous following Chol incorporation in comparison to control cells (Fig. 5D). Furthermore, the bulb width was also significantly smaller resulting in more drop-shaped caveolae (Fig. 5D’). Quantitative analysis of the spherical population of caveolae without surface connected necks allowed us to measure the surface area of caveolae. Comparison to control cells showed that area, as well as bulb width, decreased following Chol addition (Figs. 5E-E’). Furthermore, Chol incorporation resulted in a more homogeneous caveolae population in terms of size and dimensions. These data suggested that an acute increase in Chol levels in the PM of 3T3-adipocytes induced alterations in the caveolae coat architecture resulting in reduced neck diameter and a smaller more uniform bulb diameter.

**Fig. 5.**
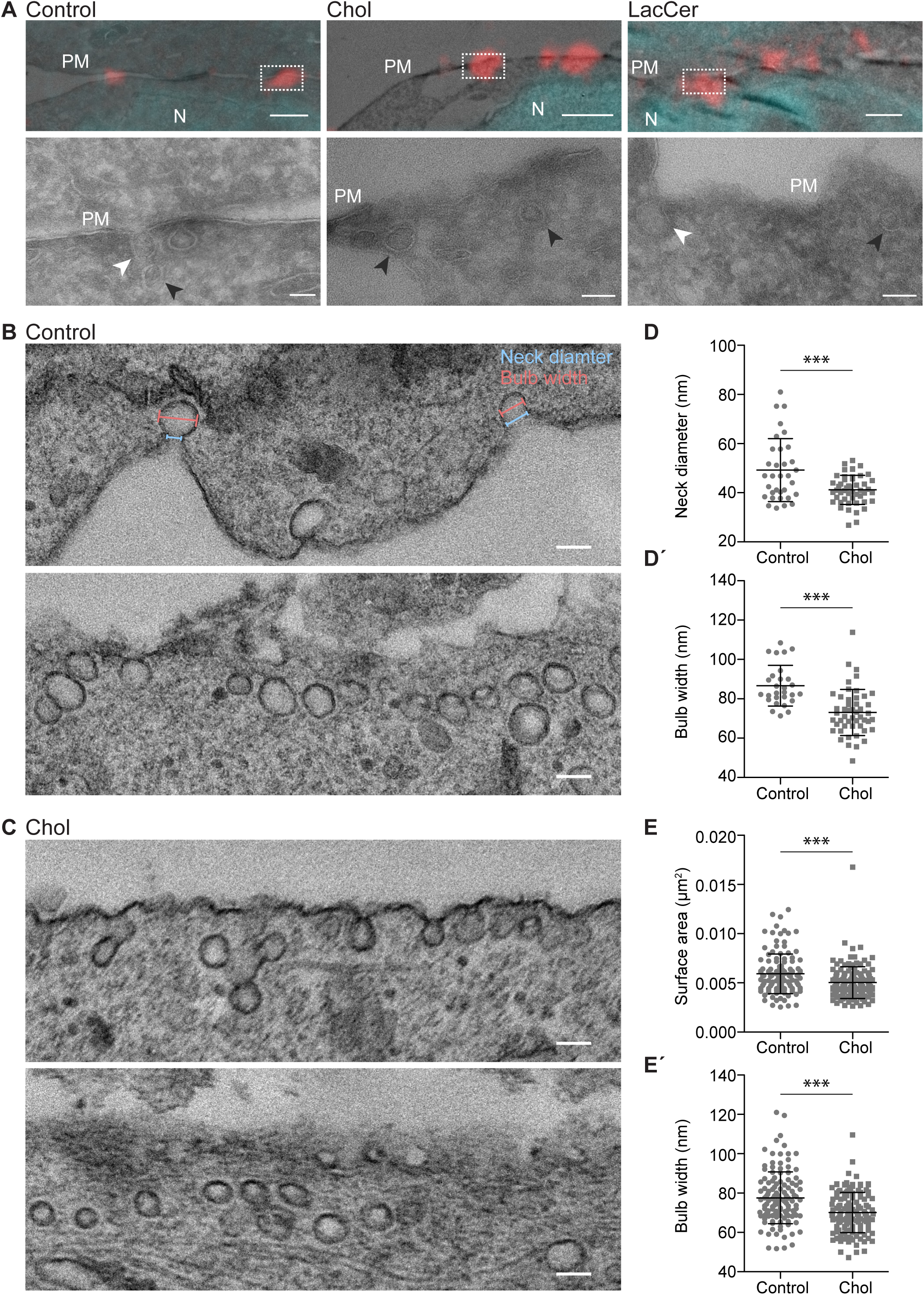
Chol accumulation reduces the caveolae diameter in 3T3 adipocytes. **(A)** Representative overlays of light microscopy images with corresponding electron micrographs showing localization of caveolae (Cav1-mCh in red) and nuclei (DAPI in cyan) for untreated Cav1-mCh HeLa cells or cells treated with Bodipy-labeled Chol or LacCer. Dotted boxes show regions of higher magnification in corresponding panels below. N, nucleus; PM, plasma membrane. White arrows denote surface connected caveolae and black arrows indicate surface adjacent caveolae. Scale bars, 1 μm; inset scale bars, 100 nm. **(B, C)** Representative electron micrographs of control 3T3-L1 adipocytes **(B)** and 3T3-L1 adipocytes treated with Bodipy-Chol **(C)**. Cells were chemically fixed, embedded in resin and processed for electron microscopy. Scale bars, 100 nm. **(D, D’)** Scatter plots showing the quantification of neck diameter **(D)** and bulb width **(D’)** of surface connected caveolae in 3T3-L1 adipocytes. Bulb width and neck diameter are highlighted in (B), upper panel. *n* ≥ 30, mean ± SEM. **(E, E’)** Scatter plots showing the quantification of surface area **(E)** and bulb width **(E’)** of surface adjacent caveolae in 3T3-L1 adipocytes. *n* ≥ 120, mean ± SEM. ***, *P* ≤ 0.001.

### GSLs are internalized to the endosomal system independent of Cav1, while Chol is predominantly trafficked to lipid droplets

Next, we aimed to address if caveolae scission significantly contributed to internalization and trafficking of lipids in our system as previously proposed (Le Lay et al., 2006; Puri et al., 2001; Shvets et al., 2015). We used fusogenic liposomes to investigate if Bodipy-labeled LacCer or Chol were internalized and trafficked through the endosomal pathway following incorporation into the PM. To mark early endosomes (EE), Rab5-BFP was transiently expressed in Cav1-mCh HeLa cells. Cells were incubated with fusogenic liposomes for either 15 min or 3 h, followed by fixation and EE localization was quantified. We observed localization of LacCer to the EE but not to the Golgi, contrasting previous studies using BSA-Bodipy-LacCer (Puri et al., 2001). After 15 min, more than half of the EE were positive for LacCer (55%) compared to only 6% for Chol (Figs. 6A-B). After 3 h, the number of LacCer-positive EE remained constant, whereas the EE positive for Chol had increased to 18% (Fig. 6B). To test if caveolae were involved in lipid trafficking to the EE, the experiments were repeated in cells depleted of Cav1 (Figs. 6C-E). After 15 min incubation time, 55% and 10% of EE were positive for LacCer and Chol, respectively (Figs. 6C-D). This suggested that while caveolae did not seem to influence the efficiency of LacCer or Chol trafficking to endosomes, loss of Cav1 resulted in an increased amount of Chol accumulating in this compartment. Our data indicate that caveolae serve as buffers or sensors of GSL and Chol concentrations rather than endocytic vesicles.

**Fig. 6.**
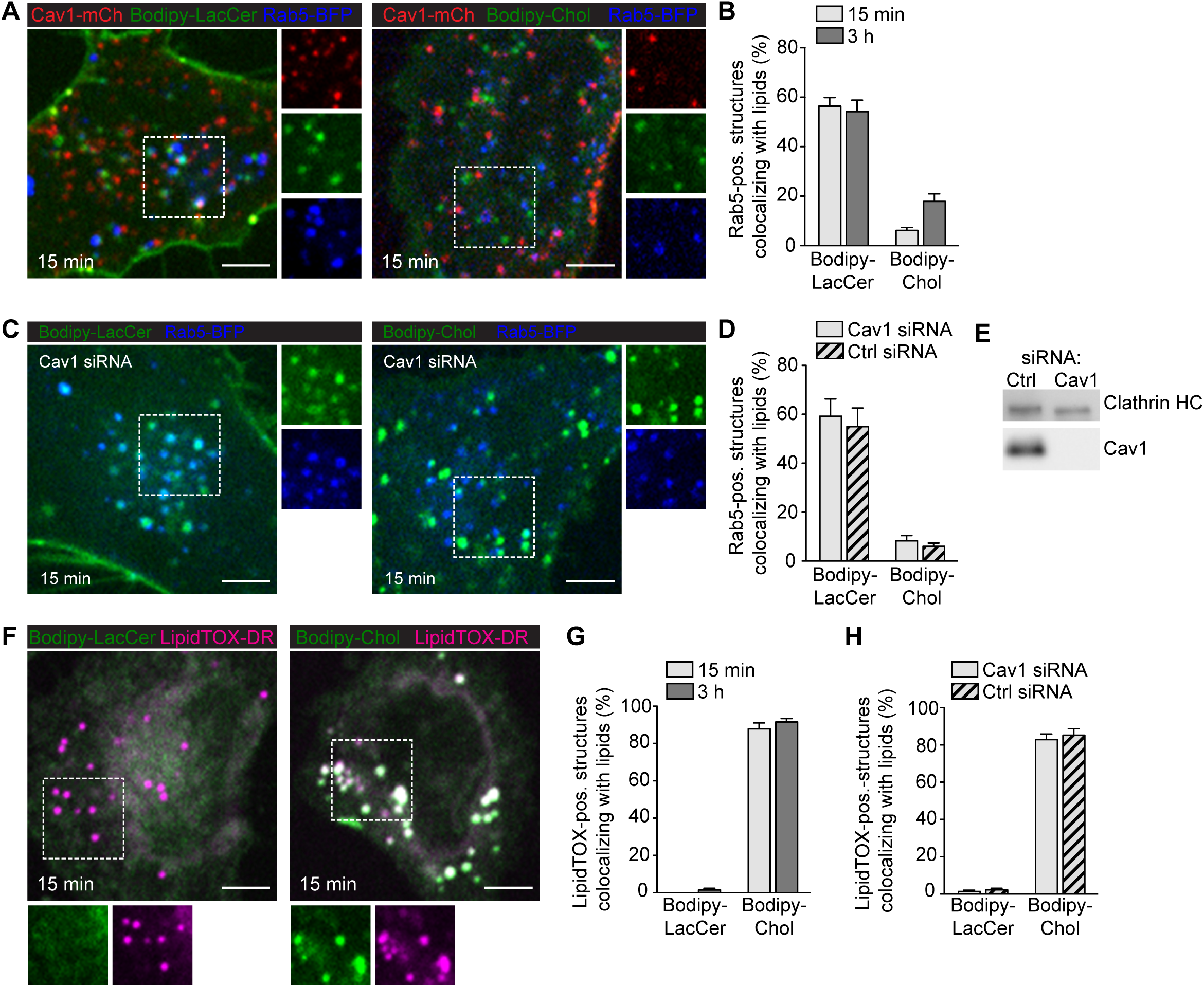
GSLs are internalized to the endosomal system independent of Cav1, while Chol is predominantly trafficked to lipid droplets. **(A)** Cav1-mCh HeLa cells expressing Rab5-BFP were incubated with Bodipy-labeled LacCer or Chol for 15 min. Individual channels are shown for selected areas (dotted box). **(B)** Colocalization of lipids with Rab5-positive structures after indicated time-points. **(C)** Cav1 siRNA-treated Cav1-mCh HeLa cells expressing Rab5-BFP after incubation with Bodipy-labeled LacCer or Chol for 15 min. High-magnification images of selected areas (dotted box) for each channel are shown. **(D)** Quantification of EE positive for lipids in cells treated with siRNA control or against Cav1. Cells were incubated with Bodipy-lipids for 15 min. **(E)** Representative immunoblots of Cav1-mCh HeLa cells treated with control siRNA or siRNA against Cav1. Clathrin HC served as loading control. **(F)** Cav1-mCh HeLa cells were incubated with Bodipy-lipids for 15 min, fixed and LDs were stained using LipidTOX-DR. **(G)** Colocalization of lipids to LDs. **(H)** Colocalization of lipids with LDs in cells depleted of Cav1 after 15 min. (B, D, E, F) *n* = 10, mean + SEM. All scale bars, 5 μm.

During our experiments, we noticed that a large fraction of Chol localized to compartments distinct from the endosomal system. To determine whether Chol localized to lipid droplets (LD) as previously proposed (Le Lay et al., 2006; Shvets et al., 2015), we incubated HeLa cells with fusogenic liposomes and visualized LD using LipidTOX-DR. On average, 85% of LDs were positive for Chol after both 15 min and 3 h (Figs. 6F-G), and similar levels of Chol-positive LD were detected in cells lacking Cav1 (Fig. 6H). While Chol extensively localized to LD, we did not observe LacCer associated with LD (Figs. 6F-G). These data are consistent with the hypothesis that excess Chol in the PM is trafficked directly to LD in a process that does not require caveolae per se, and that the levels of Chol taking an alternative route to EE increases in the absence of caveolae.

## DISCUSSION

While PM turnover is typically regulated in a tightly controlled manner, marginal changes in its composition are associated with severe diseases such as cancer, diabetes, and Alzheimer’s disease (Harayama & Riezman, 2018). It has been proposed that caveolae play a major role in preserving lipid homeostasis via sensing and buffering PM properties (Parton & del Pozo, 2013; Pilch & Liu, 2011). However, studies detailing how lipid composition influences cellular phenotypes have been hindered by a lack of methods to selectively manipulate the PM lipid composition; especially with regard to introducing specific lipids. To address this, we applied an approach for studying these systems in living cells that employs DOPE/DOTAP-based liposomes capable of mediating highly effective fusion processes with cell membranes to deliver their lipid cargoes. Such liposomes have previously been used as nanocarriers to deliver intracellular proteins (Kube et al., 2017). Our methodology successfully delivered specific lipids into the PM bilayer of living cells with high efficiency. These rapid fusion events enabled us to study, for the first time, how caveolae respond to an acute change in PM lipid composition and to observe lipid exchange in the caveolae bulb. Furthermore, the use of labelled lipids allowed us to measure the levels of incorporation in relation to endogenous levels. Our results demonstrate the power of this approach for studying caveolae dynamics and we foresee that our methodology will also be a useful tool outside of this framework.

Our work shows that the surface association of caveolae is highly sensitive to changes in the PM lipid composition. An acute increase in the levels of Chol and GSLs, which were found to specifically accumulate in caveolae, dramatically increased caveolae mobility. These caveolae traveled at higher speeds, their PM duration was shorter and they also displayed reduced levels of EHD2, a protein indicative of PM-associated caveolae. Therefore, we conclude that accumulation of Chol and GSLs in caveolae trigger surface release of caveolae. In agreement with this, analysis by EM revealed that the caveolae neck diameter was reduced in cells with elevated Chol levels. Our findings are consistent with previous reports suggesting that BSA-LacCer and Chol decrease the number of caveolae associated to the PM and enhance their mobility (Le Lay et al., 2006; Sharma et al., 2004). Based on the present study, increased caveolae mobility is a direct result of lipid accumulation in these structures. As our methodology allowed us to determine the levels of specifically incorporated lipids, we found that rapid, yet relatively small increases in specific lipids can affect caveolae dynamics. Because caveolae immediately responded to these changes in bilayer composition, we propose that they serve as PM sensors, not only for membrane tension, but also for lipid composition.

Previous studies have suggested that a threshold concentration of Chol is required to maintain caveolae integrity and proposes that assembly and disassembly is in a dynamic equilibrium dependent on Chol levels (Hailstones et al., 1998). This is also in line with our experiments showing that excess Chol drives caveolae assembly towards scission and that Chol was indeed found to accumulate in caveolae when these structures were restrained to the surface by EHD2 overexpression. Furthermore, our methodology enabled, for the first time, measurement of lipid lateral flow in and out of the caveolae bulb using FRAP. Comparing the FRAP recovery of Bodipy-LacCer and Bodipy-Chol, which were both enriched in caveolae, showed that while photobleached Bodipy-LacCer was almost fully exchanged via lateral diffusion after 2 min, photobleached Bodipy-Chol was only exchanged by 50%. This showed that Chol was sequestered in caveolae, potentially through its interaction to Cav1 (Parton & del Pozo, 2013). In contrast, Bodipy-Cer, which lacks the disaccharide structural motif of LacCer, did not accumulate in caveolae and had no effect on their dynamics, which is in agreement with earlier reports (Sharma et al., 2004). Precisely how the lactosyl group mediates the caveolae-enrichment of LacCer and how this in turn drives caveolae scission is not clear. Interestingly, we found that Bodipy-SM C_12_, but not Bodipy-SM C_5_, dramatically increased caveolae stability, in terms of both speed and duration. The contrasting effects of SM analogues were highly intriguing as both lipids are thought to partition into a liquid disordered phase in artificial giant unilamellar vesicles (Klymchenko & Kreder, 2014). However, the altered chain length might influence their interactions with Chol. Interestingly, SM has been shown to sequester a pool of Chol in the PM, which, together with an accessible and inaccessible pool, aid in the sensing of the Chol levels in the PM (Das et al., 2014). The elevated levels of SM C_12_ may alter the levels of SM-sequestered Chol, thereby affecting caveolae stability. In agreement with this, we found that a dramatic reduction of SM using SMase indeed increased the dynamics of caveolae.

While an area of extensive research, a consensus on the exact mechanism of caveolae scission has not yet been reached. Our observations suggest a model, where the accumulation of lipids in caveolae reduces the neck diameter, leading to scission. We speculate that this could be due to increased access of scission-mediating molecules like dynamin to the neck, or that that these lipids promote assembly of Cav1 and cavins, which drive curvature towards scission. The lipid-driven assembly of Cav1 may be an intrinsically unstable system, eventually resulting in scission if no restraining forces are applied. This indicates that scission is tightly coupled to, and a continuum of caveolae biogenesis. In line with this, expression of caveolin in bacterial systems induced the formation of internal caveolae-like vesicles containing caveolin so-called heterologous caveolae (Walser et al., 2012). The scission step could also involve lipid phase separation. A similar mechanism has previously been proposed but not experimentally validated (Lenz et al., 2009), and our new data shows that a locally increased concentration of GSLs and Chol in caveolae may induce phase separation and therefore facilitate budding and scission of caveolae. Consistent with this, model membrane studies have shown that sterol-induced phase separation can promote membrane scission (Bacia et al., 2005; Roux et al., 2005). Of interest, in other systems GSLs and Chol have been suggested to play a crucial role in membrane nanodomain budding to generate intracellular transport carriers (Schuck & Simons, 2004).

EHD2 has been shown to confine caveolae to the cell surface (Morén et al., 2012; Stoeber et al., 2012). In the current study, we acutely altered the lipid composition in order to induce caveolae scission and analyzed the immediate role of EHD2. We found that removal of EHD2, while at the same time changing the lipid composition, did not have an additive effect on caveolae dynamics. However, excess levels of EHD2 due to overexpression or direct microinjection, could suppress the effect of the altered lipid composition. This suggests that an increased assembly rate of EHD2 at the caveolae neck is necessary and sufficient to drive the equilibrium towards stable surface association of caveolae. We conclude that oligomers of EHD2 might provide a restraining force that prevents reduction of the neck diameter and thereby inhibits phase separation or assembly of scission-mediating proteins. Similarly, EHD2 would prevent flattening of caveolae and thus, act to stabilize the typical bulb-shape of caveolae. Therefore, EHD2 would act as a key regulator of caveolae dynamics in response to changes in both PM lipid composition and membrane tension.

Caveolae have been proposed to play an integral role in intracellular lipid trafficking (Le Lay et al., 2006; Puri et al., 2001; Shvets et al., 2015). This prompted us to examine the cellular fate of our labelled lipids. We found that while Bodipy-LacCer was internalized via the endosomal system, Chol predominately localized to LD. Importantly, and in contrast to previous data (Shvets et al., 2015), we found that loss of caveolae did not majorly influence the trafficking of these lipids in HeLa cells. Based on this, we propose that caveolae should not be considered as vehicles for internalization of lipids, but rather that lipid composition influences caveolae biogenesis and dynamics. Together with the caveolae coat components, it is feasible that sequestered lipids may control formation and define the size and curvature of these PM invaginations. This, together with our data showing that Chol is enriched and sequestered in caveolae, implies that caveolae could serve as reservoirs of Chol in the PM, thereby buffering the surface levels of this lipid.

Together, our findings indicate that the dynamic behavior of caveolae is highly sensitive to changes in PM lipid composition. We demonstrate that, following incorporation into the lipid bilayer, GSLs and Chol accumulate in caveolae, which promotes scission of these membrane invaginations from the cell surface. The current study redefines the fundamental understanding of how caveolae dynamics are governed by biologically relevant lipids and will be of future relevance linking caveolae malfunction with lipid disorders.

## MATERIALS AND METHODS

### Reagents

1,2-dioleoyl-sn-glycero-3-phosphoethanolamine (DOPE), 1,2-dioleoyl-3-trimethylammonium-propane (chloride salt) (DOTAP), TopFluor^®^-cholesterol (Bodipy-Chol), TopFluor^®^-phosphatidylethanolamine (Bodipy-PE), D-lactosyl-ß-1,1’ N-palmitoyl-D-erythro-sphingosine [LacCer(d18:1/16:0)] and Lyso-Lactosylceramide (Lyso-LacCer) were purchased from Avanti Polar Lipids Inc. (Alabaster, AL, US). Bodipy^TM^ FL C5-ganglioside GM1 (Bodipy-GM1), Bodipy^TM^ FL C5-ceramide (Bodipy-Cer), Bodipy^TM^ FL C_12_-spinghomyelin (Bodipy-SM C_12_), Bodipy^TM^ FL C_5_-spinghomyelin (Bodipy-SM C_5_) were obtained from Thermo Fisher Scientific (Waltham, MA, US). BODIPY^TM^ Fl-C5 NHS ester (4,4-Difluoro-5,7-dimethyl-4-bora-3a,4a-diaza-s-indacene-3-pentanoic acid, succinimidyl ester) was purchased from Setareh Biotech, LLC (Eugene, OR, US). Cholesterol (Chol), d7-cholesterol, *N*,*N*-diisopropylethylamine, sphingomyelinase (SMase) from *Bacillus cereus*, myriocin from *Mycelia sterilia*, anhydrous dimethylforamide (DMF), chloroform (CHCl_3_) and methanol (MeOH) were purchased from Sigma-Aldrich (St. Louis, MO, US). LC-MS grade formic acid was purchased from VWR Chemicals (Radnor, PA, US). LC-MS grade 2-propanol and acetonitrile from Merck Millipore (Billerica, MA, US). Milli-Q® water (Merck Millipore) was used. All reagents and chemicals were used without further purification.

### Bodipy-LacCer synthesis

Thin layer chromatography was performed on aluminum backed silica gel plates (median pore size 60 Å, fluorescent indicator 254 nm, Fisher Scientific, Hampton, NH, US) and visualized by exposure to UV light (365 nm) and stained with acidic ethanolic vanillin solution. Flash chromatography was performed using chromatography grade silica gel (0.035-0.070 mm, 60Å, Thermo Fisher Scientific). NMR spectra were recorded on a Bruker AVANCE (600 MHz) spectrometer. ^1^H Chemical shifts are reported in δ values relative to tetramethylsilane and referenced to the residual solvent peak (CD_3_OD: δ_H_ = 3.31 ppm, δ_C_ = 49.00 ppm). Coupling constants are reported in Hz.

Lyso-LacCer (5 mg, 8 μM) was dissolved in DMF (200 μl) and *N*,*N*-diisopropylethylamine (2.1 μl, 12 μM, 1.5 eq.) was added. BODIPY^TM^ Fl-C5 NHS ester (66 μl of a stock solution of 5 mg/100 μl DMF, 8 μM, 1.0 eq.) was added and the reaction was shielded from light and stirred for 14 h. The reaction mixture was concentrated and purified by column chromatography (CHCl_3_, MeOH, H_2_O, 70:15:2 – 65:25:2) to afford the product Bodipy-LacCer (6.5 mg, 88%, Fig. S1A) as a red film (Gretskaya & Bezuglov, 2013).

Retention factor: R_f_ = 0.46 (CHCl_3_, MeOH, H_2_O, 65:25:2)

NMR data: ^1^H-NMR (CD_3_OD, 600 MHz) δ 7.41 (1H, s), 7.03 (1H, d, *J*=4.1 Hz), 6.36 (1H, d, *J*=4.0 Hz), 6.18 (1H, s), 5.67 (1H, dt, *J*=15.3, 6.8 Hz), 5.44 (1H, dd, *J*=15.3, 7.7 Hz), 4.34 (1H, d, *J*=7.7 Hz), 4.29 (1H, d, *J*=7.8 Hz), 4.17 (1H, dd, *J*=10.1, 4.7 Hz), 4.07 (1H, t, *J*=7.9 Hz), 3.99 (1H, ddd, *J*=8.2, 4.6, 3.3 Hz), 3.89 (1H, dd, *J*=12.1, 2.6 Hz), 3.84 (1H, dd, *J*=12.2, 4.3 Hz), 3.81 (1H, d, *J*=3.1 Hz), 3.78 (1H, dd, *J*=11.4, 7.5 Hz), 3.70 (1H, dd, *J*=11.5, 4.6 Hz), 3.61 – 3.59 (1H, m), 3.62 – 3.51 (3H, m), 3.52 – 3.49 (1H, m), 3.47 (1H, dd, *J*=9.7, 3.3 Hz), 3.39 (1H, ddd, *J*=9.3, 4.0, 2.7 Hz), 3.28 (1H, t, *J*=8.5 Hz), 2.94 (2H, t, *J*=7.3 Hz), 2.50 (3H,s), 2.28 (3H, s), 2.25 (2H, t, *J*=7.0 Hz), 2.00 – 1.93 (2H, m), 1.79 – 1.66 (4H, m), 1.37 – 1.20 (31H, m, (11-CH_2_)), 0.89 (3H, t, *J*=7.0 Hz).

^13^C-NMR (CD_3_OD, 151 MHz) δ 175.7, 160.9, 160.2, 145.0, 136.1, 135.2, 134.9, 131.2, 130.0, 125.6, 120.9, 117.9, 105.1, 104.5, 80.6, 77.1, 76.5, 76.3, 74.8, 74.8, 73.0, 72.5, 70.3, 69.9, 62.5, 61.8, 54.8, 37.1, 33.4, 33.1, 30.8, 30.8, 30.8, 30.8, 30.8, 30.7, 30.5, 30.4, 30.3, 29.5, 29.4, 27.0, 23.7, 14.9, 14.5, 11.2.

### Cell lines and primary cultures

HeLa cells (ATCC-CRM-CCL-2) were cultured in Dulbecco’s Modified Eagle Medium (DMEM, Thermo Fisher Scientific) supplemented with 10% (v/v) Fetal bovine serum (FBS, Thermo Fisher Scientific) at 37°C, 5% CO2. For generation of HeLa Flp-In T-REx Caveolin1-mCherry cells the pcDNA/FRT/TO/Caveolin1-mCherry construct was generated by exchanging the EGFP-tag in the pcDNA/FRT/TO/Caveolin1-EGFP (Mohan et al., 2015) for a mCherry tag by restriction cloning using enzymes AgeI and NotI (Thermo Fisher Scientific). The HeLa Flp-In T-REx EHD2-BFP-P2A-Caveolin1-mCherry construct was generated by linearizing pcDNA/FRT/TO/Caveolin1-mCh with the restriction enzyme HindIII (Thermo Fisher Scientific). The DNA encoding EHD2-BFP and the P2A peptide was inserted by Gibson assembly using NEBuilder HiFi DNA assembly master mix (New England BioLabs, Ipswich, MA, USA). The Flp-In TRex HeLa cell lines were maintained in DMEM supplemented with 10% (v/v) FBS, 100 μg/ml hygromycin B (Thermo Fisher Scientific), and 5 μg/ml blasticidin S HCl (Thermo Fisher Scientific) for plasmid selection at 37°C, 5% CO2. Expression at endogenous levels was induced by incubation with 0.5 ng/ml (Cav1-mCh) and 1.0 ng/ml (EHD2-BFP-P2A-Cav1mCh) doxycycline hyclate (Dox, Sigma-Aldrich) for 16-24 h.

3T3-L1 fibroblasts (ATC-CL-173) were maintained in DMEM supplemented with 10% (v/v) FBS and penicillin-streptomycin (10000 U/ml, 1:100, Thermo Fisher Scientific) at 37°C, 5% CO2 and differentiated to adipocytes as previously describe (Zebisch et al., 2012). Briefly, cells were either seeded directly into a 6-well plate or on glass coverslips in a 6-well plate at 6 × 10^5^ cells/well (day −3 of differentiation). The cells reached confluency the following day and the medium was changed (day −2). After 48 h (day 0) the medium was exchanged for differentiation medium I [supplemented DMEM containing 0.5 mM 3-isobutyl-1-methylxanthine (IBMX, Sigma Aldrich), 0.25 μM dexamethasone (Dex, Sigma Aldrich), 1 μg/ml insulin (Sigma Aldrich) and 2 μM rosiglitazone (Cayman Chemical, Ann Arbor, MI, USA)]. Following incubation for 48 h, the medium was changed to differentiation medium II (supplemented DMEM containing 1 μg/ml insulin) (day 2). Experiments were performed on day 4 of differentiation.

### Fusogenic liposomes

Liposomes were prepared from a lipid mixture of DOPE, DOTAP and either Bodipy-tagged lipid or unlabeled lipid at a ratio of 47.5:47.5:5. Lipid blends were in MeOH:CHCl_3_ (1:3, v/v). Following the generation of a thin film using a stream of nitrogen gas, the vesicles were formed by addition of 20 mM HEPES (VWR, Stockholm, SE, pH 7.5, final lipid concentration 2.8 μmol/ml) and incubated for 1.5 h at room temperature. Glass beads were added to facilitate rehydration. The liposome dispersion was sonicated for 30 min (Transsonic T 310, Elma, Singen, DE). The hydrodynamic diameters (*z*-average) of the liposomes were measured using dynamic light scattering with a Malvern Zetasizer Nano-S (Malvern Instruments, Worcestershire, UK). Samples were diluted 1:100 in 20 mM HEPES (pH 7.5) and measured using a UV-transparent disposable cuvettes (Sarstedt, Nümbrecht, DE). The measurements were performed at 20°C. The Nano DTS Software 5.0 was used for acquisition and analysis of the data.

### Lipid quantification by LC-ESI-MS/MS

One day prior to experiment, cells were seeded in a 6-well plate. Cells were left untreated or treated with 11.7 nmol/ml of the different fusogenic liposomes for 10 min at 37°C, 5% CO_2_. The cells were washed three times with PBS and harvested in 500 μl MeOH by scraping. Counting revealed that approximately 4 × 10^5^ cells were obtained per sample. For myriocin (2.5 μM) and SMase (0.01 U) treatment, cells were incubated for 24 h or 2 h, respectively. Extraction was performed using a mixer mill set to a frequency 30 Hz for 2 min, with 1 tungsten carbide bead added to each tube. Thereafter the samples were centrifuged at 4°C, 14000 RPM, for 10 min. A volume of 260 μl of the supernatant was transferred to micro vials and evaporated under N_2_ (g) to dryness. The dried extracts were stored at −80°C until analysis. Calibration curves of Bodipy-labeled standards (Bodipy-SM C_12_ and Bodipy-LacCer) as well as standards for endogenous LacCer and SM [LacCer(d18:1/16:0) and SM(d18:1/16:0)] were prepared prior to analysis. Stock solutions of each compound were prepared at a concentration of 500 ng/μl and stored at −20°C. A 5-point calibration curve (0.025-0.4 ngl/μl) was prepared by serial dilutions [Bodipy-SM C_12_ R^2^ = 0.9909; LacCer(d18:1/16:0) R^2^ = 0.9945; Bodipy-LacCer R^2^ = 0.9983; LacCer(d18:1/14:0) R^2^ = 0.8742], except for endogenous SM(d18:1/16:0) where 0.025-10.0 ng/μl was used (R^2^ = 0.9991). Samples and calibration curves were analyzed using a 1290 Infinitely system from Agilent Technologies (Waldbronn, Germany), consisting of a G4220A binary pump, G1316C thermostated column compartment and G4226A autosampler with G1330B autosampler thermostat coupled to an Agilent 6490 triple quadrupole mass spectrometer equipped with a jet stream electrospray ion source operating in positive ion mode. Separation was achieved injecting 2 μl of each sample (resuspended in 20 μl of MeOH) onto a CSH C_18_ 2.1×50 mm, 1.7 μm column (Waters, Milford, MA, USA) held at 60°C in a column oven. The gradient eluents used were 60:40 acetonitrile:H_2_O (A) and 89:10.5:0.4 isopropanol:acetonitrile:water (B), both with 10 mM ammonium formate and 0.1% formic acid, with a flow rate of 500 μl/min. The initial conditions consisted of 15% B, and the following gradient was used with linear increments: 0-1.2 min (15-30% B), 1.2-1.5 (30-55% B), 1.5-4.0 (55% B), 4.0-4.8 (55-100% B), 4.8-6.8 (100% B), 7.1-8.0 (15% B). The MS parameters were optimized for each compound (Table 1). The fragmentor voltage was set at 380 V, the cell accelerator voltage at 5 V and the collision energies from 20-30 V, nitrogen was used as collision gas.

**Table 1.** Retention times (RT), MRM-transition stages monitored (precursor ion and productions) and collision energies of analyzed compounds.

| Compounds | MRM transition |  | RT (min) | Collision Energy (V) |
| --- | --- | --- | --- | --- |
|  | Precursor Ion | Product Ion |  |  |
| Bodipy-LacCer | 926.5 | 562.4 | 1.48 | 30 |
| LacCer(d18:1/16:0) | 862.6 | 520.5 | 2.84 | 20 |
| LacCer(d18:1/14:0) | 834.6 | 264.3 | 2.8 | 40 |
| SM(d18:1/16:0) | 703.6 | 184.1 | 2.9 | 30 |
| Bodipy-SM C<sub>12</sub> | 865.6 | 184.1 | 2.12 | 30 |

Jet-stream gas temperature was at 150°C with a gas flow of 16 l/min. The sheath gas temperature was kept at 350°C with a gas flow of 11 l/min. The nebulizer pressure was set to 35 psi and the capillary voltage was set at 4 kV. The QqQ was run in Dynamic MRM Mode with using a retention time delta of 0.8 min and 500 millisec cycle scans. The data was quantified using custom scripts (Swedish Metabolomics Centre, Umeå, Sweden).

### Cholesterol quantification by GC-MS

One day prior to experiment cells were seeded in a 6-well plate. Cells were left untreated or treated with 11.7 nmol/ml fusogenic liposomes for 10 min at 37°C, 5% CO_2_. The cells were washed three times with PBS and harvested in 250 μl MeOH by scraping and two wells were pooled to generate approximately 8 × 10^5^ cells per 500 μl sample into Eppendorf tubes. Extraction was performed using a mixer mill set to a frequency 30 Hz for 2 min, with 1 tungsten carbide bead added to each tube. Obtained extracts were centrifuged at 4°C, 14000 RPM for 10 min. A volume of 300 μl of the collected supernatants were transferred to individual micro vials and the extracts were dried under N_2_ (g) to dryness. Separate calibration curves were prepared for endogenous and d7-Chol. A 6-point calibration curve spanning from 0-10 ng/μl was prepared for d7-Chol (R^2^ = 0.9909). For endogenous Chol a 6-point calibration curve spanning from 0-500 ng/μl was prepared (R^2^ = 0.9969). Methyl stearate at a final concentration of 5ng/μl was used as internal standard in both calibration curves. Derivatization was performed according to a previously published method (Gullberg et al., 2004). In detail, 10 μl of methoxyamine (15 μg/μl in pyridine) was added to the dry sample that was shaken vigorously for 10 min before left to react in room temperature. After 16 hours 10 μl of MSTFA was added, the sample was shaken and left to react for 1 hour in room temperature. A volume of 10 μl of methyl stearate (15 ng/μl in heptane) was added before analysis. For d7-cholesterol quantification, 1 μl of the derivatized sample was injected by an Agilent 7693 autosampler, in splitless mode into an Agilent 7890A gas chromatograph equipped with a multimode inlet (MMI) and 10 m × 0.18 mm fused silica capillary column with a chemically bonded 0.18 μm DB 5-MS UI stationary phase (J&W Scientific). The injector temperature was 250°C. The carrier gas flow rate through the column was 1 ml min^-1^, the column temperature was held at 60°C for 1 min, then increased by 60°C min-1 to 300°C and held there for 2 min. The column effluent is introduced into the electron impact (EI) ion source of an Agilent 7000C QQQ mass spectrometer. The thermal AUX 2 (transfer line) and the ion source temperatures were 250°C and 230°C, respectively. Ions were generated by a 70 eV electron beam at an emission current of 35 μA and analyzed in dMRM-mode. The solvent delay was set to 3 min. For a list of MRM transitions see Table 2. For endogenous Chol analysis, the samples were reanalyzed in split mode (10:1) together with the Chol calibration curve. Data were processed using MassHunter Qualitative Analysis (Agilent Technologies, Atlanta, GA, USA) and custom scripts (Swedish Metabolomics Centre, Umeå, Sweden).

**Table 2.** MRM transitions for labeled and endogenous Chol.

| Compound | Comment | Precursor ion | MS1 resolution | Product ion | MS2 resolution | RT | RT delta Min (total) |
| --- | --- | --- | --- | --- | --- | --- | --- |
| **Methyl stearate** | IS-std | 298 | Unit | 101.1 | Unit | 5.6 | 2 |
| **Chol** | Quant | 329 | Unit | 95 | Unit | 7.8 | 2 |
| **Chol** | Qual | 368 | Unit | 213 | Unit | 7.8 | 2 |
| **d7-Chol** | Quant | 336 | Unit | 95 | Unit | 7.8 | 2 |
| **d7- Chol** | Qual | 375 | Unit | 213 | Unit | 7.8 | 2 |

### Calculations of the number of PM lipids

The average PM area of fibroblast is around 3000 μm^2^ (Sheetz et al., 2006), of which 23% is estimated to be occupied by proteins (Dupuy & Engelman, 2008), which translates into that the average PM of a cell contains approximately 7 × 10^9^ lipids (Alberts et al., 2002). Our data is in agreement with these reported values, whereby our measured values for SM(d18:1/16:0) being 40% of total SM species (Kjellberg et al., 2014), and 21mol% of PM lipids translating to 9.6 × 10^9^ lipids in the PM.

### Assessment of lipid incorporation into the PM with live-cell spinning disk microscopy

One day prior to the experiment, non-induced Cav1-mCh HeLa cells or 3T3-L1 adipocytes were seeded on glass coverslips (CS-25R17 or CS-25R15, Warner Instruments, Hamden, CT, US) in a 6-well plate at 3 × 10^5^ cells/well (37°C, 5% CO_2_). Live cell experiments were conducted in phenol red-free DMEM (live cell medium, Thermo Fisher Scientific) supplemented with 10% FBS and 1 mM sodium pyruvate (Thermo Fisher Scientific) at 37°C in 5% CO_2_. To follow the distribution of Bodipy throughout the PM, a POC mini 2 chamber (PeCon, Erbach, DE) was used that allowed addition of the fusogenic liposomes during data acquisition. Liposomes were added at a concentration of 7 nmol/ml and movies of confocal stacks were recorded every 30 s over a period of 5 min using a 63X lens and Zeiss Spinning Disk Confocal controlled by ZEN interface with an Axio Observer.Z1 inverted microscope, equipped with a CSU-X1A 5000 Spinning Disk Unit and an EMCCD camera iXon Ultra from ANDOR. For TIRF movies the same system was used but employing a 100X lens and an Axio Observer.Z1 inverted microscope equipped with an EMCCD camera iXonUltra from ANDOR. The increase in fluorescence intensity (FI) of the Bodipy signal was measured within circular regions of interest, which were either evenly distributed over the PM seen in the confocal section or over the basal PM in the case of TIRF. The total FI was determined by calculating integrated density (area x FI), which was then background corrected. Ten regions of interest (ROIs) per cell were analyzed using Zeiss Zen interface (*n* = 3, two independent experiments). Based on that lipids occupy 65 Å^2^, which translates to 3.1 × 10^6^ lipid molecules/μm^2^ (Dopico, 2007), and that the mean liposomes diameter was 225 nm, corresponding to an area of 0.19 μm^2^, we calculated that each liposome contained 0.6 × 10^6^ lipids, of which 5% were Bodipy-labeled. To estimate the cell volume, the cell surface was segmented with the surface feature within the Imaris x64 9.1.2 (Bitplane, Zurich, CH) using the mCherry fluorescence.

### Constructs, transfections and cell treatments

pTagBFP-C (Evrogen, Moscow, RU) was used to generate the expression constructs of Rab5 and EHD2 wt or I157Q. Cav1-mCh HeLa cells were transfected with Lipofectamine^TM^ 2000 (Thermo Fisher Scientific) using Opti-MEM™ I reduced serum medium (Thermo Fisher Scientific) for transient protein expression. For EHD2 and Cav1 depletion, Cav1-mCh HeLa cells were transfected with either stealth siRNA, specific against human EHD2 or human Cav1, or scrambled control (all from Thermo Fisher Scientific) using Lipofectamine^TM^ 2000 and Opti-MEM according to manufacturer’s instructions unless otherwise stated. Cells were transfected twice over a period of 72 h before the experiment. Protein levels were analyzed by SDS-PAGE and immunoblotting using rabbit anti-EHD2 (Morén et al., 2012) and rabbit anti-Cav1 antibodies (Abcam, Cambridge, UK). Mouse anti-clathrin heavy chain (clone 23, BD Transduction Laboratories, San Jose, CA, US) was used as loading control. Cells were treated with 2.5 μM myriocin in complete medium 24 h prior to harvesting. SMase was added to cells to generate a final concentration of 0.01 units in complete medium 2 h prior to harvesting or live cell imaging.

### Analysis of caveolae dynamics

To track caveolae dynamics, induced Cav1-mCh HeLa cells were treated with fusogenic liposomes (7 nmol/ml) and 5 min TIRF movies were recorded with an acquisition time of 3 s. Imaris software was used for tracking analysis of Cav1-mCh positive structures, which were segmented as spots and structures with a diameter of 0.4 μm were selected. The applied algorithm was based on Brownian motion with max distance travelled of 0.8 μm and a max gap size of 4. Experiments where EHD2 (wt and mutant) was either transiently expressed or depleted were performed and analyzed the same way. Colocalization of EHD2 (wt and mutant) to Cav1-mCh was quantified with Imaris software. Within a ROI, spots were created in one channel (*e.g.*, red channel) and the second channel (*e.g.*, blue channel) was masked. The masked spots show only colocalized red and blue spots and the percentage was correlated to the original channel. The analysis of the dynamic behavior of caveolae positive for or lacking EHD2-BFP was preformed using double Flp-In EHD2-BFP Cav1-mCh HeLa cells. The tracking was done as described above and the data from the tracks of Cav1-mCh spots lacking EHD2-BFP was collected and removed from the data of Cav1-mCh spots positive for EHD2-BFP. Statistical analysis was performed on track duration (s) and track mean speed (μm/s) data and data is shown as fold change. All micrographs and acquired movies were prepared with Fiji (Schindelin et al., 2012) and Adobe Photoshop CS6.

### Intracellular trafficking of lipids

Induced Cav1-mCh HeLa cells were seeded on glass coverslips (CS-25R15) in a 6-well plate at 3 × 10^5^ cells/well (37°C, 5% CO_2_). On the following day, the cells were incubated with fusogenic liposomes (7 nmol/ml) for 15 min or 3 h. Rab5-BFP (Francis et al., 2015) was either transiently expressed or. To analyze the localization of lipids to lipid droplets (LDs), induced Cav1-mCh HeLa cells were treated with lipids for 15 min or 3 h, fixed and stained with HCS LipidTOX™ Deep Red Neutral Lipid Stain (1:200, Thermo Fisher Scientific). Confocal stacks were acquired on Zeiss Spinning Disk Confocal microscope. The colocalization was analyzed as described above. Micrographs were prepared with Fiji (Schindelin et al., 2012) and Adobe Photoshop CS6.

### Immunostaining

Induced Cav1-mCh HeLa cells were seeded on precision coverslips (No. 1.5H, Paul Marienfeld GmbH & Co. KG, Lauda-Königshofen, DE) in 24-well plates at 50 × 10^3^ cells/well and incubated overnight (37°C, 5% CO_2_). Following incubation with fusogenic liposomes (7 nmol/ml) for 1 h, the cells were washed thrice with phosphate-buffered saline (PBS, pH 7.4). Cells were fixed with 4 % PFA in PBS (Electron Microscopy Sciences, Hatfield, PA, US) and subsequent permeabilization and blocking was carried out simultaneously using PBS containing 5% goat serum and 0.05% saponin. Cells were then immunostained with rabbit anti-EHD2 (Morén et al., 2012) and rabbit anti-PTRF (Abcam) followed by goat anti-rabbit IgG secondary antibody coupled to Alexa Fluor 647 (Thermo Fisher Scientific) as previously described (Lundmark et al., 2008). Confocal images were acquired using the Zeiss Spinning Disk Confocal microscope (63X lens). Pearson colocalization coefficients were obtained using Imaris software applying the Coloc feature with automatic thresholding. All Pearson coefficients were derived from two independent experiments for the EHD2 stain. Analysis of the colocalization of cavin 1 and Cav1-mCh was repeated once. Data from at least 30 images was analyzed with images containing 2–3 cells on average. Micrographs were prepared using Fiji (Schindelin et al., 2012) and Adobe Photoshop CS6.

### FRAP experiments

Induced Cav1-mCh HeLa cells were seeded on glass coverslips (CS-25R15) in a 6-well plate at 3 × 10^5^ cells/well and incubated overnight (37°C, 5% CO_2_). Cells were treated with 7 nmol/ml of Bodipy-labeled liposomes for 10 min followed by two washes with live cell media before imaging using TIRF using a Zeiss Axio Observer.Z1 inverted microscope. Three reference images were recorded before a ROI was photobleached for 1000 ms using maximal laser intensity (488 nm or 561 nm). The fluorescent recovery images were taken every 3 s for 5 min. For the lipid incorporation experiment, a region within the PM with homogeneous fluorescence was chosen. FRAP of the EHD2 mutants was performed the same way. For the LacCer and Chol accumulated in caveolae, FRAP was preformed between 15 to 60 min after lipid addition and regions with structures positive for Cav1-mCh, EHD2-BFP and Bodipy-lipid were selected. For FRAP experiments that quantified the recovery of Cav1-mCh, induced Cav1-mCh HeLa cells were either untreated, depleted of EHD2 using siRNA or incubated with Bodipy-LacCer liposomes. FRAP experiments were performed as described above using the Zeiss Spinning Disk Confocal microscope (63X lens). The signal recovery monitored in focal plane close to the basal membrane. The intensities of the bleached regions were corrected for background signal and photobleaching of the cell. Data from at least 10 cells were collected per condition and mean FRAP recovery curves were plotted using Prism 5.0 (GraphPad, San Diego, CA, US).

### Microinjection

Mouse EHD2 cysteine mutant construct (L303C,C96S, C138S, C356S) was expressed as N-terminal His_6_-tag fusion proteins in *Escherichia coli* Rosetta (DE3) and purified (Daumke et al., 2007). Dithiothreitol was removed from the protein using PD-10 columns and the protein was labelled with Alexa Fluor™ 647 C2 Maleimide (Thermo Fisher Scientific) (Hoernke et al., 2017). The protein was diluted to a concentration of 0.5 mg/ml in 150 mM NaCl, 20 mM HEPES pH 7.5 and 1 mM MgCl_2_. Cav1-mCh HeLa cells were transfected with siRNA and induced as described above. One day prior to the injection experiment, Cav1-mCh HeLa cells were seeded in MatTek dishes (35 mm dish, high tolerance 1.5, MatTek Corporation, Ashland, MA, US) with a cell density of 3 × 10^5^ cells/dish and induced with Dox. In the case of LacCer addition, the cells were treated with 7 nmol/ml of Bodipy-LacCer fusogenic liposomes for 10 min followed by two washes with live cell media before microinjection. Microinjection was preformed with Injectman NI2 coupled to the programmable microinjector Femtojet (Eppendorf, Hamburg, DE). The protein was loaded in Femtotips II (Eppendorf) and injection was done with an injection pressure of 1.0 hPa, compensation pressure of 0.5 hPa and injection time of 0.1 s. Live images were acquired on TIRF every 3 s for a total of five min using a Nikon Eclipse Ti-E inverted microscope with a 100X lens (Apochromat 1.49 Oil 0.13-0.20 DIC N2, Nikon). Z-stacks of injected cells were captured using a 60X lens (Apochromat 1.4o Oil DIC, Nikon). Tracking of Cav1-mCh and colocalization analysis was done with Imaris as previously described.

### Correlative light electron microscopy

Cav1-mCh cells transiently expressing EHD2-GFP alone or treated with either Bodipy-LacCer or Bodipy-Chol liposomes were fixed in 2% paraformaldehyde (PFA) and 0.2% of glutaraldehyde (Taab Laboratory Equipment Ltd, Aldermaston, UK) in 0.1 M phosphate buffer (pH 7.4) for 1-2 h and then stored in 1% PFA at 4°C. For the grid preparation, the cells were scraped into the fixative solution and washed thrice with PBS (pH 7.4) and once with PBS containing 0.1% glycine (pH 7.4, Merck Millipore, Burlington, US). The cell pellet was embedded in 12% gelatin (Dr. Oetker, food grade) in 0.1 M phosphate buffer (pH 7.4). Blocks of around 1 mm^2^ were cut and cryo-protected by overnight infiltration in 2.3 M sucrose (VWR) in 0.1 M phosphate buffer. Next, the blocks were plunge frozen in liquid nitrogen. The sample block was sectioned at −120°C to obtain 80 nm sections. These were mounted in a drop of in 0.1 M phosphate buffer containing 1:1 of 2% methyl cellulose (Sigma-Aldrich) and 2.3 M sucrose on TEM grids with a carbon-coated Formvar film (Taab Laboratory Equipment Ltd). The grids were incubated with PBS (pH 7.4) at 37°C for 20 min and stained with DAPI (4’,6-Diamidino-2-Phenylindole, Dilactate, 1:1000 in PBS, pH 7.4, Thermo Fisher Scientific) before imaging on a Nikon Eclipse Ti-E inverted microscope with a 100X lens (Apochromat 1.49 Oil 0.13-0.20 DIC N2, Nikon). Low magnification images were taken at 20X for orientation on the grid and to aid the overlay of fluorescent microscopy images and with the higher resolution images of TEM. Contrasting for TEM was done by embedding the grids in 1.8% methyl cellulose and 0.4% uranyl acetate (Polysciences, Inc., Hirschberg an der Bergstrasse, DE) solution prepared in water (pH 4) for 10 min in the dark. TEM was performed with a Talos 120C transmission electron microscope (FEI, Eindhoven, NL) operating at 120kV. Micrographs were acquired with a Ceta 16M CCD camera (FEI) using Maps 3.3 (FEI, Hillsboro, OR, US). The fluorescent images were overlaid atop TEM images of the same cells collected from the ultrathin section using Adobe Photoshop CS6.

### Electron microscopy

3T3-L1 cells were seeded on MatTek dishes (35 mm dish, high tolerance 1.5) and differentiated to adipocytes as described above. 3T3-L1 adipocytes were untreated or incubated with Bodipy-Chol liposomes for 45 min, washed with PBS and fixed as follows.

All chemical fixation steps were performed using a microwave (Biowave, TED PELLA, inc.) unless stated and solutions were prepared and rinses were performed in 0.1M cacodylate buffer (Sigma-Aldrich) or water. Fixation of the cells was performed in 0.05% malachite green oxalate (Sigma-Aldrich) and 2.5% gluteraldehyde (Taab Laboratory Equipment Ltd, Aldermaston, UK) in cacodylate buffer. The samples were rinsed four times with cacodylate buffer and post-fixed with 0.8% K_3_Fe(CN)_6_ (Sigma-Aldrich) and 1% OsO_4_ (Sigma-Aldrich) in cacodylate buffer and rinsed four times with cacodylate buffer. The samples were then stained with 1% aqueous tannic acid (Sigma-Aldrich). Following two rinses in cacodylate buffer and water, samples were stained with 1% aqueous uranyl acetate (Polysciences, Inc., Hirschberg an der Bergstrasse, DE). After four washed with water, samples were dehydrated in gradients of ethanol (25%, 50%, 75%, 90%, 95%, 100% and 100%) (VWR). The samples were infiltrated with graded series of hard grade spurr resin (Taab Laboratory Equipment Ltd, Aldermaston, UK) in ethanol (1:3, 1:1 and 3:1) and then left in 100% resin for 1 h at room temperature. The samples were later polymerized overnight at 60°, sectioned and imaged with a Talos 120C transmission electron microscope (FEI, Eindhoven, NL) operating at 120kV. To obtain quantitative data, segmentation of caveolae for measurement of bulb width and measurement of neck diameter for surface-connected caveolae was performed with “icy” (de Chaumont et al., 2012). In order to extract bulb width and surface area, “active cells” plug-in was used with three points to make an elliptical contour that fitted individual caveolae. The neck diameter was obtained by drawing a ROI across the neck of surface-connected caveolae. The analysis was performed blinded and with randomized sections.

### Statistical analysis

Statistical analysis was carried out by two-tailed unpaired Student *t*-test for comparison with control samples using GraphPad Prism 5.0 software. All experiments were performed at least twice with data representing mean ± SEM unless otherwise stated.

## ONLINE SUPPLEMENTAL MATERIAL

Fig. S1 illustrates dynamic behavior of caveolae at the cell surface and shows characterization of liposomes as well as chromatography data. Fig. S2 shows the correlation between track duration and track mean speed and the effect of GSLs and Chol on lifetime of caveolae. Fig. S3 shows the colocalization of EHD2 to Cav1 in double Flp-In cells, analysis of track duration in presence of EHD2-I157Q-BFP and TIRF images of TIRF images of Cav1-mCh HeLa cells with or without microinjection of EHD2-647. Fig. S4 shows that Bodipy-labeled LacCer and Chol accumulate in caveolae at the PM and demonstrates the recovery of lipids within caveolae after photobleaching. Fig. S5 shows CLEM approach and upregulation of EHD2 and Cav1 in 3T3-L1 adipocytes. Video 1-4 show cell surface dynamics of Cav1-mCh untreated (Video 1) or following addition of Bodipy-LacCer (Video 2), Chol (Video 3) or Bodipy-SM C_12_ (Video 4). Video 5-6 show colocalization of Bodipy-LacCer (Video 5) or Bodipy-Chol (Video 6) with Cav1-mCh positive structures. Video 7-8 show accumulation of Bodipy-LacCer (Video 7) or Chol (Video 8) in Cav1-mCh positive structures in presence of EHD2-BFP.

## ACKNOWLEDGMENTS

We acknowledge the Biochemical Imaging Center (BICU) and Umeå Core Facility Electron Microscopy (UCEM) at Umeå University and the National Microscopy Infrastructure, NMI (VR-RFI 2016-00968) for providing assistance. We especially thank Irene Martinez at BICU for assistance and expertise with image analysis and data visualization. We thank Mikkel Roland Holst for help with establishing the HeLa Flp-In T-REx Caveolin1-mCherry cells. This work was supported by the Swedish Cancer Society (CAN2014/746, CAN 2017/735, R.L. and M.H.), The Hagbergs Foundation (R.L. and M.H.), Kempe Foundation (L.W.K.M.) and the Swedish Research Council (dnr 2017-04028, R.L. and E.L.).

## AUTHOR CONTRIBUTIONS

M.H., E.L. and R.L. designed the research; M.H., E.L., N.G.V.G. and R.L. performed research and analyzed data; L.W.K.M. synthesized lipids, M.A. and A.J. performed mass spectrometry, M.H., E.L., L.W.K.M. and R.L. wrote the paper.

## Supplemental Material

**Fig. S1.**
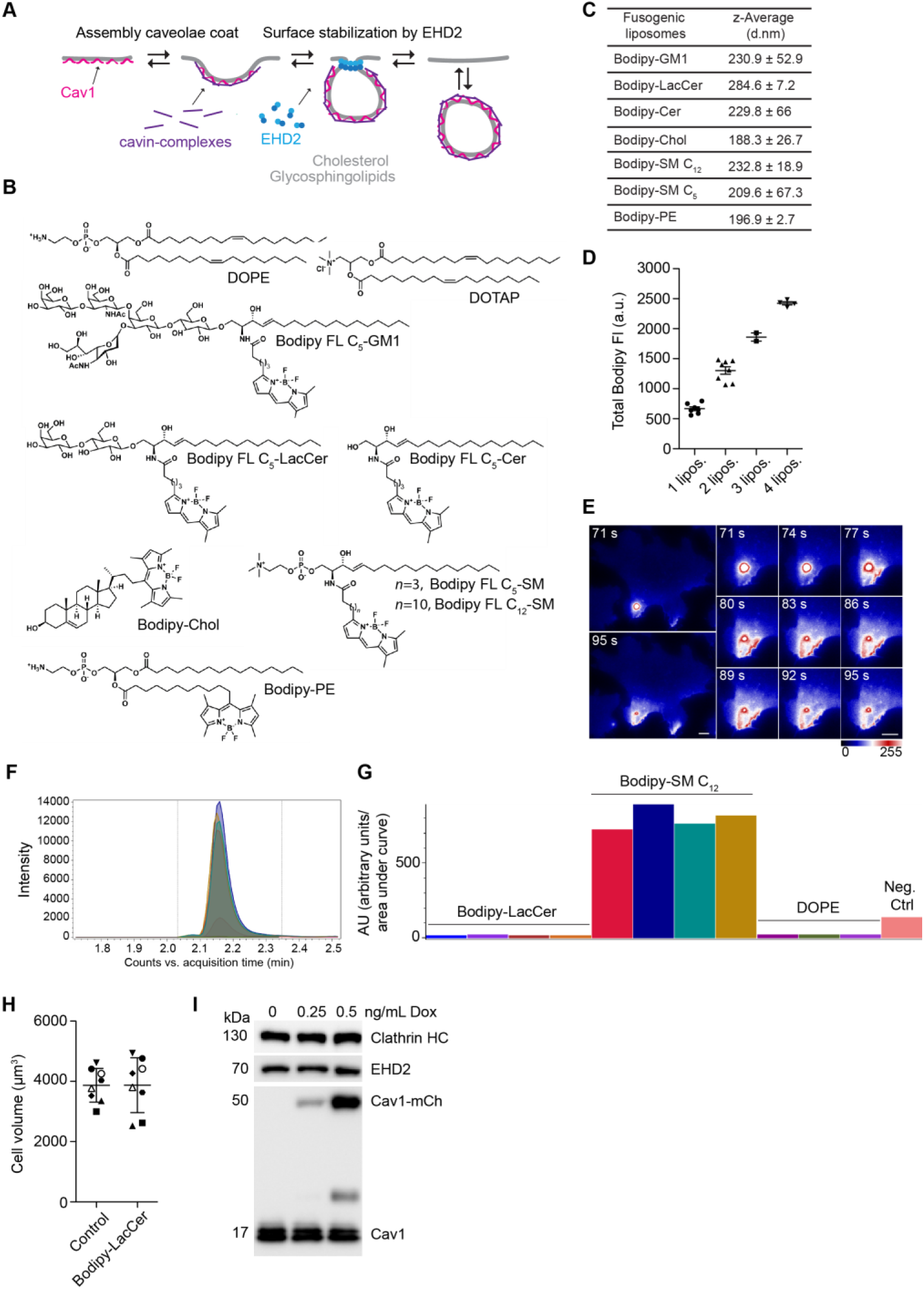
Liposome characterization and incorporation efficiencies of Bodipy-lipids measured by mass spectrometry. **(A)** Scheme illustrating caveolae dynamics at PM. Caveolae formation and coat assembly are primarily driven by the integral membrane protein Cav1 and cavin proteins. EHD2 controls surface association of caveolae. **(B)** Chemical structures of lipids used in this study. **(C)** Hydrodynamic diameter as *z*-average of DOPE:DOTAP:Bodipy-lipid liposomes. *n* = 3, three independent experiments, mean ± SD. **(D)** Total Bodipy FI of liposomes containing Bodipy-LacCer was determined in a single confocal section (0.5 μm) using spinning disk microscopy. Bodipy FI corresponds to the number of liposomes measured in each ROI. *n* = 22, mean ± SEM. **(E)** Time-lapse imaging of a vesicle fusing with the PM. A single fusion event highlights the rapid distribution of the fluorophore from the liposome-membrane contact site and subsequent fusion Distribution of Bodipy fluorescence is intensity coded using LUT. Scale bars, 10 μm. **(F, G)** Visualization of chromatography of Bodipy-SM C_12_. Samples visualized: Bodipy-LacCer treated samples, Bodipy-SM C_12_ treated samples, DOPE control samples and a negative control of Bodipy-SM C_12_ (liposomes added to wells without cells). **(F)** Chromatography of the samples. **(G)** Integrated area of each individual sample. **(H)** Analysis of the cell volume before and after addition of Bodipy-LacCer liposomes. Cell surface was segmented with Imaris using mCh fluorescence. Identical symbols in control and Bodipy-LacCer represent the same cell. *n* = 8, mean ± SD. **(I)** Representative immunoblots showing protein expression of EHD2, Cav1-mCh and Cav1 after induction of Cav1-mCh HeLa cells with different concentrations of Dox. Clathrin HC served as loading control. Related to Fig. 1.

**Fig. S2.**
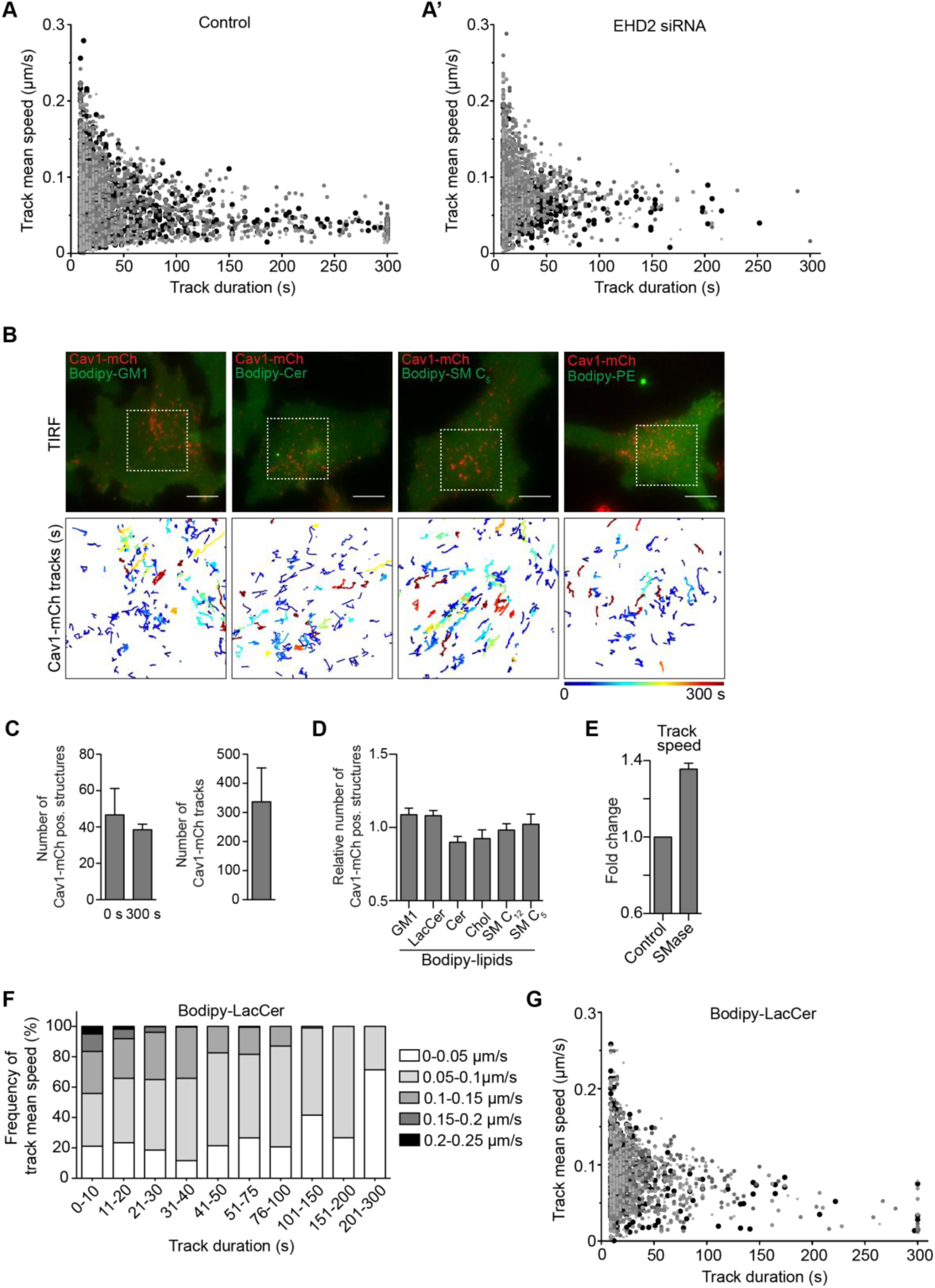
GSLs and Chol affect lifetime of caveolae while caveolae numbers at the PM are unchanged. **(A, A’)** Correlation between track duration and track mean speed. TIRF live cell movies of Cav1-mCh structures **(A)** and cells lacking EHD2 **(A’)** were analyzed (five datasets for each condition). Identical symbols represent tracks from the same cell. **(B)** Representative images from TIRF live cell movies of Dox-induced Cav1-mCh HeLa cells after incubation with different fusogenic liposomes containing Bodipy-lipids (final total lipid concentration of 7 nmol/mL) for 15 min. Cav1-mCh structures were tracked using Imaris software. Color-coded trajectories illustrate time that structures can be tracked at PM over 5 min (dotted square). Scale bars, 10 μm. **(C)** Number of Cav1-mCh positive structures at the beginning and at the end of 5 min TIRF movies and the corresponding number of tracks detected. *n* ≥ 8, three independent experiments, mean + SEM. **(D)** Relative number of caveolae at the PM of Cav1-mCh HeLa cells before and after addition of fusogenic liposomes. TIRF live cell movies from Fig. 2C and S2B were analyzed. Number of caveolae after lipid treatment was normalized to the number of caveolae in control cells. *n* ≥ 8, three independent experiments, mean + SEM. **(E)** Quantification of track mean speed of Cav1-mCh structures from TIRF movies following incubation with SMase for 2 h. Fold changes are relative to control (Cav1-mCh). *n* ≥ 5, mean + SEM. **(F)** Distribution of track mean speed in subpopulations of track duration of Cav1-mCh structures treated with Bodipy-LacCer liposomes. **(G)** Correlation between track duration and track mean speed of Cav1-mCh structures treated with Bodipy-LacCer liposomes. In (F) and (G) five datasets were analyzed. All analysis was performed using Imaris software. Related to Fig. 2.

**Fig. S3.**
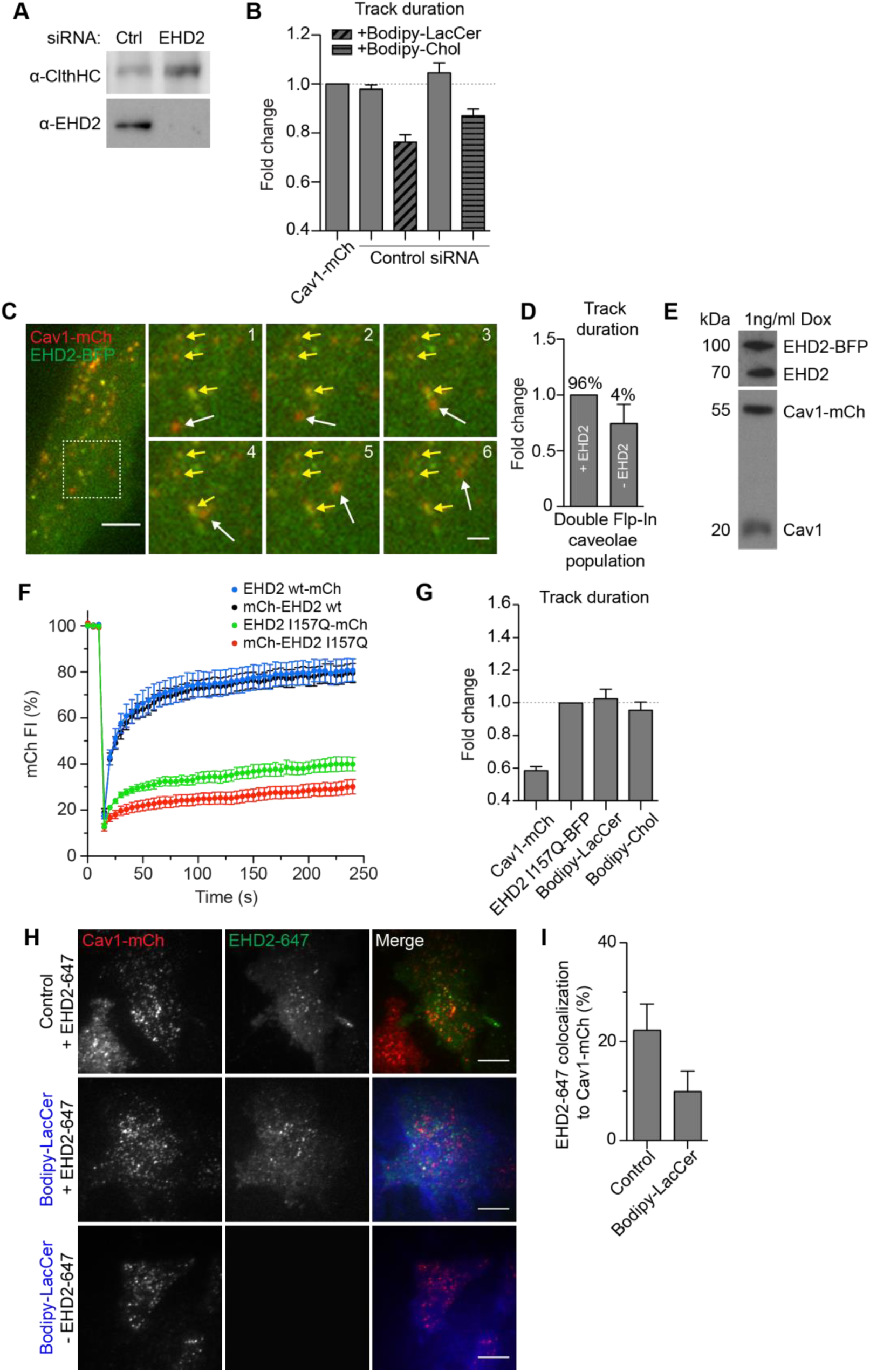
Stabilization of caveolae to the PM by EHD2 and EHD2-I157Q cannot be reversed by addition of Bodipy-labeled LacCer or Chol. **(A)** Representative immunoblots of Cav1-mCh HeLa cells treated with Ctrl siRNA or siRNA against EHD2. Clathrin HC served as loading control. **(B)** Effect of lipids on track duration of Cav1-mCh structures analyzed following control siRNA-treatment. *n* ≥ 8, two independent experiments, mean + SEM. **(C)** Representative time-lapse images of Cav1-mCh positive for EHD2-BFP (yellow arrows) or lacking EHD2-BFP (white arrows) in double Flp-In EHD2-BFP Cav1-mCh HeLa cells. Dotted box shows higher magnification region. Numbering corresponds to number of frames. Scale bar, 10 μm; inset scale bars, 2 μm. **(D)** Differences in track duration of Cav1-mCh structures positive for EHD2-BFP or lacking EHD2-BFP in double Flp-In EHD2-BFP Cav1-mCh HeLa cells. Percentage of Cav1-mCh structures positive or lacking EHD2-BFP are indicated. *n* = 8, mean + SEM. **(E)** Representative immunoblots of double Flp-In EHD2-BFP Cav1-mCh HeLa cells induced with 1 ng/ml Dox. **(F)** FRAP curves of mCh-tagged EHD2 wt or EHD2 I157Q expressing HeLa cells. A ROI was photobleached and recovery of mCherry fluorescence intensity (mCherry FI) was monitored. Intensities were normalized to background and reference. *n* = 8, mean ± SEM. **(G)** Cav1-mCh HeLa cells transiently expressing EHD2-I157Q-BFP were incubated with Bodipy-LacCer or Bodipy-Chol liposomes and track duration was analyzed. *n* ≥ 8, two independent experiments, mean + SEM. **(H)** Representative live cell TIRF images of Cav1-mCh HeLa cells untreated or treated with Bodipy-LacCer and with or without microinjection of EHD2-647. **(I)** Quantification of the colocalization of microinjected EHD2-647 to Cav1-mCh in control cells and cells treated with Bodipy-LacCer liposomes prior to injection. *n* ≥ 5, mean + SEM. Scale bar, 10 μm. In (B, D, G) Imaris software was used to analyze data. Changes in track duration are relative to control (indicated by dotted line). Related to Fig. 3.

**Fig. S4.**
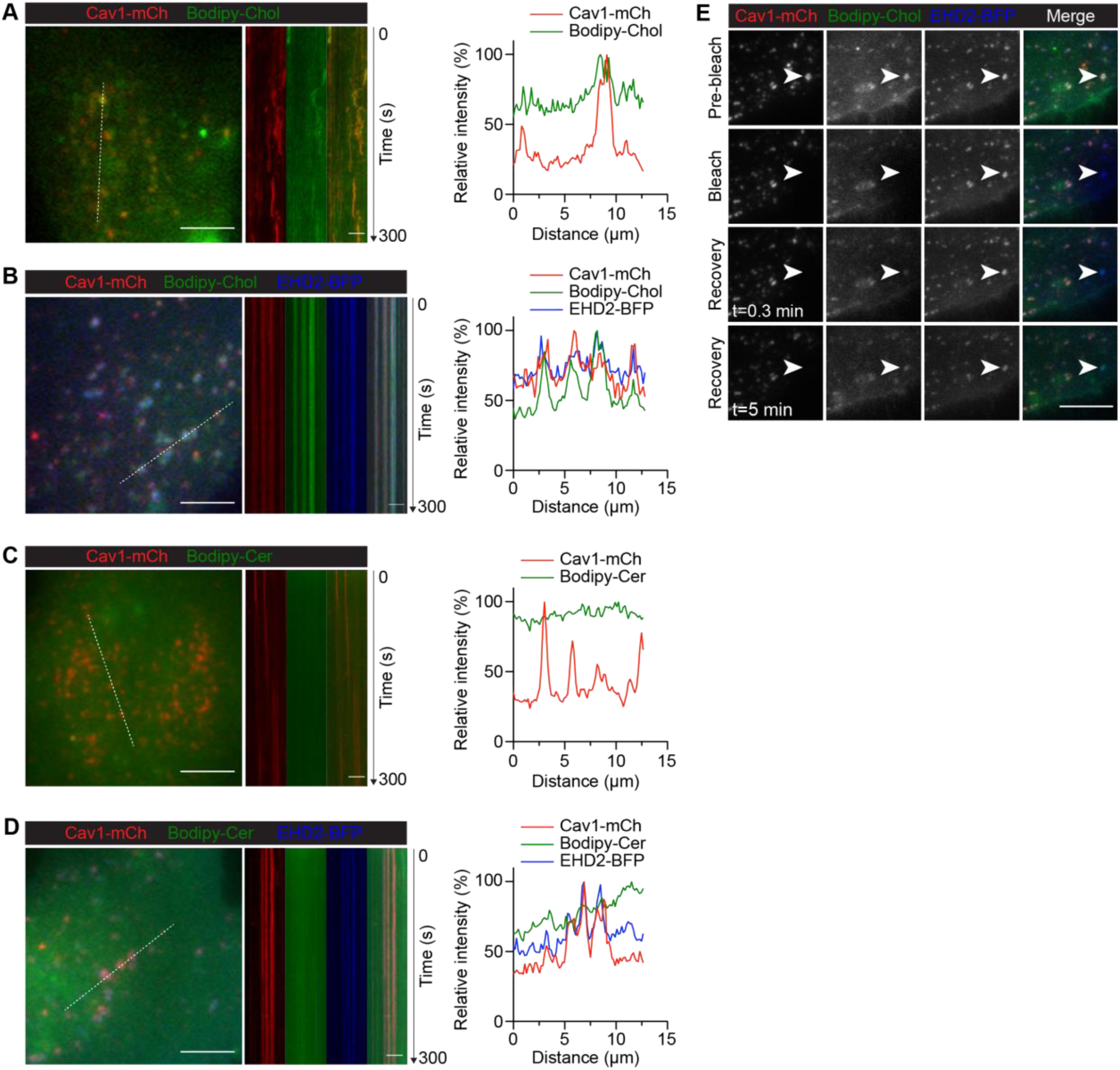
LacCer and Chol but not Cer accumulate in caveolae at the PM and recover within caveolae following photobleaching. **(A, B)** Cav1-mCh HeLa cells **(A)** and Cav1-mCh HeLa cells transiently expressing EHD2-BFP **(B)** were incubated with Bodipy-Chol liposomes. White lines indicate the location of the kymograph and the corresponding intensity profiles illustrate the colocalization of Bodipy-Chol with Cav1-mCh either alone or in the presence of EHD2-BFP. Intensity profiles are relative to the maximum value for each sample. **(C, D)** As for (A, B), cells were incubated with Bodipy-Cer. Scale bars, 10 μm; kymograph scale bars, 5 μm. **(E)** Bodipy fluorescence recovery experiments to study the accumulation of lipids in caveolae. Cav1-mCh HeLa cells transiently expressing EHD2-BFP were incubated with Bodipy-Chol liposomes for 10 min. Following photobleaching, the fluorescence recovery of the Bodipy signal within caveolae was monitored over time. White arrow highlights surface connected caveolae with accumulated Chol. Scale bars, 5 μm. Related to Fig. 4.

**Fig. S5.**
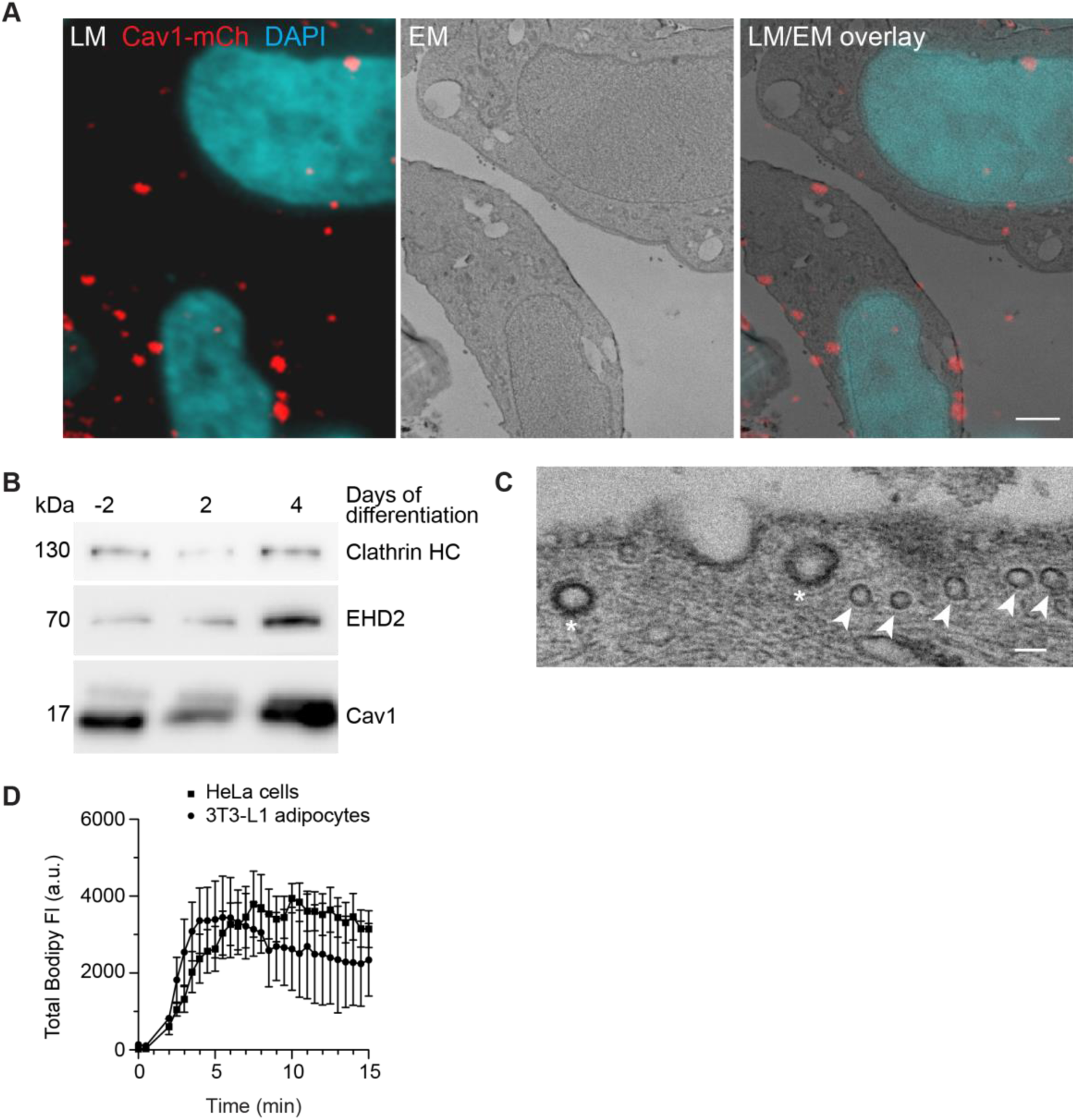
Expression of EHD2 and Cav1 is upregulated in 3T3-L1 adipocytes. **(A)** Cav1-mCh HeLa cells were induced with Dox and transiently expressed EHD2-GFP. Light microscopy image showing localization of caveolae (Cav1-mCh) and nuclei (DAPI) within cells (LM, left panel). Middle panel depicts corresponding EM images. Overlay of LM and EM images shows correlation of fluorescently labeled structures to ultrastructure in same cells (right panel). Scale bar, 2 μm. **(B)** Representative immunoblots showing protein expression of EDH2 and Cav1 during 3T3-L1 differentiation. Clathrin HC served as loading control. **(C)** Representative electron micrographs of 3T3-L1 adipocytes. Caveolae (indicated by arrow) can be clearly distinguished from clathrin-coated pits (indicated by asterisk). Scale bar, 100 nm. **(D)** Incorporation rate of Bodipy-Chol into the PM of live 3T3-L1 adipocytes. Cells were treated with fusogenic liposomes (final total lipid concentration 7 nmol/mL). Total fluorescence intensity (FI) of the Bodipy signal was measured within circular ROIs in a confocal section using spinning disk microscopy. Ten ROIs were analyzed using the Zeiss Zen system software. *n* = 2, mean ± SEM. Related to Fig. 5.

**Video 1. Cell surface dynamics of Cav1-mCh.** A representative TIRF live cell movie of Dox-induced Cav1-mCh HeLa cells. The image in Fig. 2C (Cav1-mCh) is taken from this movie. Movie in real time spans 5 min and was recorded at 3 s intervals. Scale bar, 10 μm. Related to Fig. 2.

**Video 2. Cell surface dynamics of Cav1-mCh after treatment with Bodipy-LacCer.** A representative TIRF live cell movie of Dox-induced Cav1-mCh HeLa cells after 15 min incubation with liposomes containing Bodipy-LacCer. The image in Fig. 2C is taken from this movie. Movie in real time spans 5 min and was recorded at 3 s intervals. Scale bar, 10 μm. Related to Fig. 2.

**Video 3. Cell surface dynamics of Cav1-mCh after treatment with Bodipy-Chol.** A representative TIRF live cell movie of Dox-induced Cav1-mCh HeLa cells after 15 min incubation with liposomes containing Bodipy-Chol. The image in Fig. 2C is taken from this movie. Movie in real time spans 5 min and was recorded at 3 s intervals. Scale bar, 10 μm. Related to Fig. 2.

**Video 4. Cell surface dynamics of Cav1-mCh after treatment with Bodipy-SM C_12_.** A representative TIRF live cell movie of Dox-induced Cav1-mCh HeLa cells after 15 min incubation with liposomes containing Bodipy-SM C_12_. The image in Fig. 2C is taken from this movie. Movie in real time spans 5 min and was recorded at 3 s intervals. Scale bar, 10 μm. Related to Fig. 2.

**Video 5. Bodipy-LacCer colocalizes with Cav1-mCh positive structures.** A representative TIRF live cell movie of Dox-induced Cav1-mCh HeLa cells after incubation with liposomes containing Bodipy-LacCer. The image in Fig. 4A is taken from this movie and corresponds to the ROI highlighted by the white square. Movie in real time spans 5 min and was recorded at 3 s intervals. Scale bar, 10 μm. Related to Fig. 4.

**Video 6. Bodipy-Chol colocalizes with Cav1-mCh positive structures.** A representative TIRF live cell movie of Dox-induced Cav1-mCh HeLa cells after incubation with liposomes containing Bodipy-Chol. The image in Fig. S4A is taken from this movie and corresponds to the ROI highlighted by the white square. Movie in real time spans 5 min and was recorded at 3 s intervals. Scale bar, 10 μm. Related to Fig. 4.

**Video 7. Bodipy-LacCer accumulates in caveolae.** A representative TIRF live cell movie of Dox-induced Cav1-mCh HeLa cells transiently expressing EHD2-BFP after incubation with liposomes containing Bodipy-LacCer. The image in Fig. 4B is taken from this movie and corresponds to the ROI highlighted by the white square. Movie in real time spans 5 min and was recorded at 3 s intervals. Scale bar, 10 μm. Related to Fig. 4.

**Video 8. Bodipy-Chol accumulates in caveolae.** A representative TIRF live cell movie of Dox-induced Cav1-mCh HeLa cells transiently expressing EHD2-BFP after incubation with liposomes containing Bodipy-Chol. The image in Fig. S4B is taken from this movie and corresponds to the ROI highlighted by the white square. Movie in real time spans 5 min and was recorded at 3 s intervals. Scale bar, 10 μm. Related to Fig. 4.

## REFERENCES

Alberts, B., Johnson, A., Lewis, J., Raff, M., Roberts, K., & Walter, P. (2002). Molecular Biology of the Cell (4 ed.). New York: Garland Science.

Bacia, K., Schwille, P., & Kurzchalia, T. (2005). Sterol structure determines the separation of phases and the curvature of the liquid-ordered phase in model membranes. Proc. Natl. Acad. Sci. USA, 102(9), 3272–3277. doi:10.1073/pnas.0408215102

Boucrot, E., Howes, M. T., Kirchhausen, T., & Parton, R. G. (2011). Redistribution of caveolae during mitosis. J. Cell Biol., 124(12), 1965–1972. doi:10.1242/jcs.076570

Cao, H., Alston, L., Ruschman, J., & Hegele, R. A. (2008). Heterozygous CAV1 frameshift mutations (MIM 601047) in patients with atypical partial lipodystrophy and hypertriglyceridemia. Lipids Health Dis., 7(1), 3. doi:10.1186/1476-511x-7-3

Cohen, A. W., Hnasko, R., Schubert, W., & Lisanti, M. P. (2004). Role of Caveolae and Caveolins in Health and Disease. Physiol. Rev., 84(4), 1341–1379. doi:10.1152/physrev.00046.2003

Crescencio, M. E., Rodríguez, E., Páez, A., Masso, F. A., Montaño, L. F., & López-Marure, R. (2009). Statins inhibit the proliferation and induce cell death of human papilloma virus positive and negative cervical cancer cells. J. Biomed. Sci., 5(4), 411–420.

Csiszar, A., Hersch, N., Dieluweit, S., Biehl, R., Merkel, R., & Hoffmann, B. (2010). Novel fusogenic liposomes for fluorescent cell labeling and membrane modification. Bioconjug. Chem., 21(3), 537–543. doi:10.1021/bc900470y

Das, A., Brown, M. S., Anderson, D. D., Goldstein, J. L., & Radhakrishnan, A. (2014). Three pools of plasma membrane cholesterol and their relation to cholesterol homeostasis. Elife, 3, e02882. doi:10.7554/eLife.02882

Daumke, O., Lundmark, R., Vallis, Y., Martens, S., Butler, P. J. G., & McMahon, H. T. (2007). Architectural and mechanistic insights into an EHD ATPase involved in membrane remodelling. Nature, 449(7164), 923–927. doi:10.1038/nature06173

de Chaumont, F., Dallongeville, S., Chenouard, N., Hervé, N., Pop, S., Provoost, T., Meas-Yedid, V., Pankajakshan, P., Lecomte, T., Le Montagner, Y., Lagache, T., Dufour, A., & Olivo-Marin, J.-C. (2012). Icy: an open bioimage informatics platform for extended reproducible research. Nat. Meth., 9, 690. doi:10.1038/nmeth.2075

Dopico, A. M. (2007). Methods in Membrane Lipids (1 ed.). New York City: Humana Press.

Dupuy, A. D., & Engelman, D. M. (2008). Protein area occupancy at the center of the red blood cell membrane. Proc Natl Acad Sci USA, 105(8), 2848–2852. doi:10.1073/pnas.0712379105

Francis, M. K., Holst, M. R., Vidal-Quadras, M., Henriksson, S., Santarella-Mellwig, R., Sandblad, L., & Lundmark, R. (2015). Endocytic membrane turnover at the leading edge is driven by a transient interaction between Cdc42 and GRAF1. J Cell Sci, 128(22), 4183–4195. doi:10.1242/jcs.174417

Gretskaya, N. M., & Bezuglov, V. V. (2013). Synthesis of BODIPY® FL C5-Labeled D-erythro- and L-threo-Lactosylceramides. Chem Nat Compd, 49(1), 17–20. doi:10.1007/s10600-013-0494-3

Gullberg, J., Jonsson, P., Nordstrom, A., Sjostrom, M., & Moritz, T. (2004). Design of experiments: an efficient strategy to identify factors influencing extraction and derivatization of Arabidopsis thaliana samples in metabolomic studies with gas chromatography/mass spectrometry. Anal Biochem, 331(2), 283–295. doi:10.1016/j.ab.2004.04.037

Gulshan, K., Brubaker, G., Wang, S., Hazen, S. L., & Smith, J. D. (2013). Sphingomyelin Depletion Impairs Anionic Phospholipid Inward Translocation and Induces Cholesterol Efflux. Journal of Biological Chemistry. doi:10.1074/jbc.M113.512244

Hailstones, D., Sleer, L. S., Parton, R. G., & Stanley, K. K. (1998). Regulation of caveolin and caveolae by cholesterol in MDCK cells. J Lipid Res, 39(2), 369–379.

Harayama, T., & Riezman, H. (2018). Understanding the diversity of membrane lipid composition. Nat. Rev. Mol. Cell Biol., 19, 281. doi:10.1038/nrm.2017.138

Hayashi, Y. K., Matsuda, C., Ogawa, M., Goto, K., Tominaga, K., Mitsuhashi, S., Park, Y. E., Nonaka, I., Hino-Fukuyo, N., Haginoya, K., Sugano, H., & Nishino, I. (2009). Human PTRF mutations cause secondary deficiency of caveolins resulting in muscular dystrophy with generalized lipodystrophy. J. Clin. Invest., 119(9), 2623–2633. doi:10.1172/jci38660

Hirama, T., Das, R., Yang, Y., Ferguson, C., Won, A., Yip, C. M., Kay, J. G., Grinstein, S., Parton, R. G., & Fairn, G. D. (2017). Phosphatidylserine dictates the assembly and dynamics of caveolae in the plasma membrane. J. Biol. Chem., 292(34), 14292–14307. doi:10.1074/jbc.M117.791400

Hoernke, M., Mohan, J., Larsson, E., Blomberg, J., Kahra, D., Westenhoff, S., Schwieger, C., & Lundmark, R. (2017). EHD2 restrains dynamics of caveolae by an ATP-dependent, membrane-bound, open conformation. Proc. Natl. Acad. Sci. USA, 114(22), E4360–E4369. doi:10.1073/pnas.1614066114

Kim, C. A., Delepine, M., Boutet, E., El Mourabit, H., Le Lay, S., Meier, M., Nemani, M., Bridel, E., Leite, C. C., Bertola, D. R., Semple, R. K., O’Rahilly, S., Dugail, I., Capeau, J., Lathrop, M., & Magre, J. (2008). Association of a homozygous nonsense caveolin-1 mutation with Berardinelli-Seip congenital lipodystrophy. J. Clin. Endocrinol. Metab., 93(4), 1129–1134. doi:10.1210/jc.2007-1328

Kjellberg, M. A., Backman, A. P. E., Ohvo-Rekilä, H., & Mattjus, P. (2014). Alternation in the Glycolipid Transfer Protein Expression Causes Changes in the Cellular Lipidome. PLoS ONE, 9(5), e97263. doi:10.1371/journal.pone.0097263

Kleusch, C., Hersch, N., Hoffmann, B., Merkel, R., & Csiszár, A. (2012). Fluorescent Lipids: Functional Parts of Fusogenic Liposomes and Tools for Cell Membrane Labeling and Visualization. Molecules, 17(1), 1055. doi:10.3390/molecules17011055

Klymchenko, Andrey S., & Kreder, R. (2014). Fluorescent Probes for Lipid Rafts: From Model Membranes to Living Cells. Cell Chem. Biol., 21(1), 97–113. doi:10.1016/j.chembiol.2013.11.009

Krause, B. R., & Hartman, A. D. (1984). Adipose tissue and cholesterol metabolism. J. Lipid Res., 25(2), 97–110.

Kube, S., Hersch, N., Naumovska, E., Gensch, T., Hendriks, J., Franzen, A., Landvogt, L., Siebrasse, J.-P., Kubitscheck, U., Hoffmann, B., Merkel, R., & Csiszár, A. (2017). Fusogenic Liposomes as Nanocarriers for the Delivery of Intracellular Proteins. Langmuir, 33(4), 1051–1059. doi:10.1021/acs.langmuir.6b04304

Le Lay, S., Hajduch, E., Lindsay, M. R., Le Lièpvre, X., Thiele, C., Ferré, P., Parton, R. G., Kurzchalia, T., Simons, K., & Dugail, I. (2006). Cholesterol-Induced Caveolin Targeting to Lipid Droplets in Adipocytes: A Role for Caveolar Endocytosis. Traffic, 7(5), 549–561. doi:10.1111/j.1600-0854.2006.00406.x

Lenz, M., Morlot, S., & Roux, A. (2009). Mechanical requirements for membrane fission: Common facts from various examples. FEBS Lett., 583(23), 3839–3846. doi:10.1016/j.febslet.2009.11.012

Liu, L., Brown, D., McKee, M., Lebrasseur, N. K., Yang, D., Albrecht, K. H., Ravid, K., & Pilch, P. F. (2008). Deletion of Cavin/PTRF causes global loss of caveolae, dyslipidemia, and glucose intolerance. Cell Metab., 8(4), 310–317. doi:10.1016/j.cmet.2008.07.008

Lorizate, M., Sachsenheimer, T., Glass, B., Habermann, A., Gerl, M. J., Kräusslich, H.-G., & Brügger, B. (2013). Comparative lipidomics analysis of HIV-1 particles and their producer cell membrane in different cell lines. Cellular Microbiology, 15(2), 292–304. doi:10.1111/cmi.12101

Lundmark, R., Doherty, G. J., Howes, M. T., Cortese, K., Vallis, Y., Parton, R. G., & McMahon, H. T. (2008). The GTPase-Activating Protein GRAF1 Regulates the CLIC/GEEC Endocytic Pathway. Curr. Biol., 18(22-2), 1802–1808. doi:10.1016/j.cub.2008.10.044

Matthäus, C., Lahmann, I., Kunz, S., Jonas, W., Melo, A. A., Lehmann, M., Larsson, E., Lundmark, R., Kern, M., Blüher, M., Müller, D. N., Haucke, V., Schürmann, A., Birchmeier, C., & Daumke, O. (2019). EHD2-mediated restriction of caveolar dynamics regulates cellular lipid uptake. bioRxiv, 511709. doi:10.1101/511709

Mohan, J., Morén, B., Larsson, E., Holst, M. R., & Lundmark, R. (2015). Cavin3 interacts with cavin1 and caveolin1 to increase surface dynamics of caveolae. J. Cell Biol., 128(5), 979–991. doi:10.1242/jcs.161463

Morén, B., Hansson, B., Negoita, F., Fryklund, C., Lundmark, R., Göransson, O., & Stenkula, K. G. (2019). EHD2 regulates adipocyte function and is enriched at cell surface– associated lipid droplets in primary human adipocytes. Mol. Biol. Cell, 30(10), 1147–1159. doi:10.1091/mbc.E18-10-0680

Morén, B., Shah, C., Howes, M. T., Schieber, N. L., McMahon, H. T., Parton, R. G., Daumke, O., & Lundmark, R. (2012). EHD2 regulates caveolar dynamics via ATP-driven targeting and oligomerization. Mol. Biol. Cell, 23(7), 1316–1329. doi:10.1091/mbc.E11-09-0787

Nassoy, P., & Lamaze, C. (2012). Stressing caveolae new role in cell mechanics. Trends Cell Biol., 22(7), 381–389. doi:10.1016/j.tcb.2012.04.007

Örtegren, U., Karlsson, M., Blazic, N., Blomqvist, M., Nystrom, F. H., Gustavsson, J., Fredman, P., & Stralfors, P. (2004). Lipids and glycosphingolipids in caveolae and surrounding plasma membrane of primary rat adipocytes. Eur. J. Biochem., 271(10), 2028–2036. doi:10.1111/j.1432-1033.2004.04117.x

Parton, R. G., & del Pozo, M. A. (2013). Caveolae as plasma membrane sensors, protectors and organizers. Nat. Rev. Mol. Cell Biol., 14(2), 98–112. doi:10.1038/nrm3512

Pelkmans, L., & Zerial, M. (2005). Kinase-regulated quantal assemblies and kiss-and-run recycling of caveolae. Nature, 436, 128. doi:10.1038/nature03866

Pilch, P. F., & Liu, L. (2011). Fat caves: caveolae, lipid trafficking and lipid metabolism in adipocytes. Trends Endocrinol. Metab., 22(8), 318–324. doi:10.1016/j.tem.2011.04.001

Puri, V., Watanabe, R., Singh, R. D., Dominguez, M., Brown, J. C., Wheatley, C. L., Marks, D. L., & Pagano, R. E. (2001). Clathrin-dependent and -independent internalization of plasma membrane sphingolipids initiates two Golgi targeting pathways. J. Cell Biol., 154(3), 535–548. doi:10.1083/jcb.200102084

Razani, B., Combs, T. P., Wang, X. B., Frank, P. G., Park, D. S., Russell, R. G., Li, M., Tang, B., Jelicks, L. A., Scherer, P. E., & Lisanti, M. P. (2002). Caveolin-1-deficient Mice Are Lean, Resistant to Diet-induced Obesity, and Show Hypertriglyceridemia with Adipocyte Abnormalities. J. Biol. Chem., 277(10), 8635–8647. doi:10.1074/jbc.M110970200

Rothberg, K. G., Heuser, J. E., Donzell, W. C., Ying, Y.-S., Glenney, J. R., & Anderson, R. G. W. (1992). Caveolin, a protein component of caveolae membrane coats. Cell, 68(4), 673–682. doi:10.1016/0092-8674(92)90143-Z

Roux, A., Cuvelier, D., Nassoy, P., Prost, J., Bassereau, P., & Goud, B. (2005). Role of curvature and phase transition in lipid sorting and fission of membrane tubules. EMBO J., 24(8), 1537–1545. doi:10.1038/sj.emboj.7600631

Schindelin, J., Arganda-Carreras, I., Frise, E., Kaynig, V., Longair, M., Pietzsch, T., Preibisch, S., Rueden, C., Saalfeld, S., Schmid, B., Tinevez, J.-Y., White, D. J., Hartenstein, V., Eliceiri, K., Tomancak, P., & Cardona, A. (2012). Fiji: an open-source platform for biological-image analysis. Nat. Meth., 9(7), 676–682. doi:10.1038/nmeth.2019

Schuck, S., & Simons, K. (2004). Polarized sorting in epithelial cells: raft clustering and the biogenesis of the apical membrane. J. Cell Biol., 117(25), 5955–5964. doi:10.1242/jcs.01596

Sharma, D. K., Brown, J. C., Choudhury, A., Peterson, T. E., Holicky, E., Marks, D. L., Simari, R., Parton, R. G., & Pagano, R. E. (2004). Selective Stimulation of Caveolar Endocytosis by Glycosphingolipids and Cholesterol. Mol. Biol. Cell, 15(7), 3114–3122. doi:10.1091/mbc.E04-03-0189

Sheetz, M. P., Sable, J. E., & Dobereiner, H. G. (2006). Continuous membrane-cytoskeleton adhesion requires continuous accommodation to lipid and cytoskeleton dynamics. Annu. Rev. Biophys. Biomol. Struct., 35, 417–434. doi:10.1146/annurev.biophys.35.040405.102017

Shvets, E., Bitsikas, V., Howard, G., Hansen, C. G., & Nichols, B. J. (2015). Dynamic caveolae exclude bulk membrane proteins and are required for sorting of excess glycosphingolipids. Nat. Commun., 6. doi:10.1038/ncomms7867

Singh, R. D., Liu, Y., Wheatley, C. L., Holicky, E. L., Makino, A., Marks, D. L., Kobayashi, T., Subramaniam, G., Bittman, R., & Pagano, R. E. (2006). Caveolar Endocytosis and Microdomain Association of a Glycosphingolipid Analog Is Dependent on Its Sphingosine Stereochemistry. J. Biol. Chem., 281(41), 30660–30668. doi:10.1074/jbc.M606194200

Singh, R. D., Marks, D. L., Holicky, E. L., Wheatley, C. L., Kaptzan, T., Sato, S. B., Kobayashi, T., Ling, K., & Pagano, R. E. (2010). Gangliosides and beta1-integrin are required for caveolae and membrane domains. Traffic 11(3), 348–360. doi:10.1111/j.1600-0854.2009.01022.x

Singh, R. D., Puri, V., Valiyaveettil, J. T., Marks, D. L., Bittman, R., & Pagano, R. E. (2003). Selective Caveolin-1–dependent Endocytosis of Glycosphingolipids. Mol. Biol. Cell, 14(8), 3254–3265. doi:10.1091/mbc.E02-12-0809

Sinha, B., Köster, D., Ruez, R., Gonnord, P., Bastiani, M., Abankwa, D., Stan, R. V., Butler-Browne, G., Vedie, B., Johannes, L., Morone, N., Parton, R. G., Raposo, G., Sens, P., Lamaze, C., & Nassoy, P. (2011). Cells Respond to Mechanical Stress by Rapid Disassembly of Caveolae. Cell, 144(3), 402–413. doi:10.1016/j.cell.2010.12.031

Stoeber, M., Stoeck, I. K., Hänni, C., Bleck, C. K. E., Balistreri, G., & Helenius, A. (2012). Oligomers of the ATPase EHD2 confine caveolae to the plasma membrane through association with actin. EMBO J., 31(10), 2350–2364. doi:10.1038/emboj.2012.98

Thorn, H., Stenkula, K. G., Karlsson, M., Ortegren, U., Nystrom, F. H., Gustavsson, J., & Stralfors, P. (2003). Cell surface orifices of caveolae and localization of caveolin to the necks of caveolae in adipocytes. Mol. Biol. Cell, 14(10), 3967–3976. doi:10.1091/mbc.e03-01-0050

Walser, Piers J., Ariotti, N., Howes, M., Ferguson, C., Webb, R., Schwudke, D., Leneva, N., Cho, K.-J., Cooper, L., Rae, J., Floetenmeyer, M., Oorschot, Viola M. J., Skoglund, U., Simons, K., Hancock, John F., & Parton, Robert G. (2012). Constitutive Formation of Caveolae in a Bacterium. Cell, 150(4), 752–763. doi:10.1016/j.cell.2012.06.042

Zebisch, K., Voigt, V., Wabitsch, M., & Brandsch, M. (2012). Protocol for effective differentiation of 3T3-L1 cells to adipocytes. Anal. Biochem., 425(1), 88–90. doi:10.1016/j.ab.2012.03.005

